# Basophils drive the resolution and promote wound healing in adult and aged mice

**DOI:** 10.1101/2024.07.02.601665

**Authors:** Julie Bex, Victoria Peter, Chaimae Saji, Léa Chapart, Suoqin Jin, Gregory Gautier, Olivier Thibaudeau, Morgane K. Thaminy, John Tchen, Quentin Simon, Xing Dai, Kensuke Miyake, Hajime Karasuyama, Jean X. Jiang, Karmella Naidoo, Marc Benhamou, Ulrich Blank, Renato Monteiro, Graham Le Gros, Nicolas Charles, Christophe Pellefigues

## Abstract

An active resolution is critical to control the duration of inflammation and limit its pathological consequences. Defects in resolution during wound healing allow the emergence of chronic wounds, a common complication in the elderly. Here, we show that basophils infiltrate the periphery of mouse skin wounds for at least three weeks, during both the inflammation and resolution phases of wound healing. Depletion of basophils induces an increased secretion of inflammatory molecules, accumulation and activation of pro-inflammatory leukocytes, and delays the wound healing response. Basophils particularly promote epidermal differentiation towards homeostasis in the wounds. Basophil-derived IL-4 and M-CSF drive partly their immunoregulatory and healing properties. Unexpectedly, aged mice basophils infiltrate more potently the wounds to promote inflammation resolution, showing a transcriptomic signature biased towards tissue remodeling. Thus, basophils are not pro-inflammatory but pro-resolution cells during an essential biological process such as skin wound healing. Unraveling basophil pro-resolution properties may reveal new strategies to fight chronic wounds in the elderly.

**SUMMARY:** The resolution of inflammation is essential for homeostasis and becomes defective with age. Here we show basophil immunoregulatory properties promote resolution during skin wound healing, in both adult and aged mice.

## Introduction

The delicate balance between pro-inflammatory, anti-inflammatory, and pro-resolution mechanisms during an inflammatory response needs to be coordinated to avoid excessive amplitude, duration, and pathological consequences in response to danger or damage. Uncontrolled inflammation can promote cytokine storms, fibrosis, or chronic wounds, especially in an increasingly fragile proportion of the population suffering from chronic inflammation such as the elderly. Immune cells are important actors in initiating, amplifying, or resolving inflammation during both infectious and sterile physiological responses(Furman *et al*., 2019).

Basophils are rare circulating “type 2” granulocytes involved in allergic or autoimmune diseases and immunity against various parasites, known for their pro-inflammatory effector functions, and their pro-Th2 or Th17 immunomodulatory functions. However, basophils manifest both pro-inflammatory and pro-resolving properties in models of chronic skin allergic inflammation(Egawa *et al*., 2013) or acute atopic dermatitis(Pellefigues, Naidoo, *et al*., 2021). They can control the extent or amplitude of inflammation through various mechanisms(Miyake, Ito and Karasuyama, 2022; Poto *et al*., 2023), including extracellular ATP degradation(Tsai *et al*., 2015) and secretion of anti-inflammatory mediators such as retinoic acid(Hachem *et al*., 2023), IL-10(Kleiner *et al*., 2021), or the type 2 cytokines IL-4 and IL-13 (Pellefigues, Mehta, *et al*., 2021). Basophils are the main source of IL-4 in models of helminth infection(van Panhuys *et al*., 2011), atopic dermatitis(Pellefigues, Naidoo, *et al*., 2021; Leyva-Castillo *et al*., 2022; Takahashi *et al*., 2023) or skin infection(Leyva-Castillo *et al*., 2021). In *vivo*, basophil-derived IL-4 is sufficient to dampen epidermal type 17 inflammation against infections(Leyva-Castillo *et al*., 2021). It also fosters the emergence of immunosuppressive myeloid cells in the bone marrow(LaMarche *et al*., 2024) and promotes an M2-like macrophage bias and the resolution of inflammation in various experimental models of chronic skin allergic or atopic inflammation(Egawa *et al*., 2013; Pellefigues, Naidoo, *et al*., 2021; Miyake *et al*., 2024), liver infection(Blériot *et al*., 2015) and heart infarction(Sicklinger *et al*., 2021). Type 2 cytokines particularly promote monocyte-to-resident macrophage transition(Finlay *et al*., 2023), local macrophage proliferation(Jenkins *et al*., 2011) and protect macrophages from immunosenescence to dampen age-associated inflammation and frailty(Zhou *et al*., 2024). Basophils also secrete M-CSF, a growth factor critical for macrophage homeostasis. Basophil-derived M-CSF seems important for alveolar macrophage development(Cohen *et al*., 2018) and promotes the resolution of skin atopic inflammation(Pellefigues, Naidoo, *et al*., 2021). M-CSF works alongside IL-4 to regulate resident macrophage homeostasis in *vivo*(Jenkins *et al*., 2013). In the skin, basophils promote keratinocyte differentiation and homeostasis in a model of epidermal inflammation(Strakosha *et al*., 2024) and during atopic-like inflammation(Pellefigues, Naidoo, *et al*., 2021). As basophils express a very high diversity of ligands to communicate with both immune and non-hematopoietic cells(Cohen *et al*., 2018; Cui *et al*., 2024), they represent unique players to fine-tune the resolution of inflammatory events.

Here, we studied the role of basophils during both the inflammation and resolution phases in mouse models of wound healing. We found that basophils quickly infiltrated the periphery of skin wounds, and persisted for at least three weeks. During both the inflammatory and resolution phases, constitutive or conditional depletion of basophils led to an accumulation of pro-inflammatory mediators and immune cell activation, delayed differentiation of wound keratinocytes and wound closure. Basophil-derived IL-4 and M-CSF showed complex immunoregulatory properties in the wounds but IL-4 was critical for a monocyte to macrophage transition during the resolution phase. As resolution becomes defective with aging, we also analyzed the pro-resolution capabilities of basophils in aged mice. Unexpectedly, aged mice showed an increased basophil infiltration, which kept their pro-resolving capabilities. This was confirmed by the reanalysis of a recently published scRNAseq dataset collected on mouse back skin wounds (Vu *et al*., 2022). Overall, basophils drive the resolution of inflammation and accelerate skin wound healing, and these properties remain potent upon aging.

## Results

### 1) Basophils infiltrate skin wounds early during inflammation and are activated during the resolution phase coinciding with the emergence of M2-like macrophages

We used 2mm circular sterile punch biopsies to generate incisional wounds in mouse ears (“Ear punch” model) and monitored the kinetics of the local skin leukocyte infiltrate by flow cytometry. We observed an early significant peak of leukocyte accumulation at 24h, followed by their progressive disappearance during the three weeks of the study. Consensual pro-inflammatory cells such as neutrophils, Ly6C+ monocytes, and Ly6C+ macrophages exhibited similar kinetics and their proportions strongly decreased at one week post wound. More than 90% of macrophages expressed Ly6C on Day 1. Another population of monocytes expressing CD16.2 (coded by *Fcgr4*), and eosinophils, did not show any enrichment during the course of the study. We also observed an early accumulation of basophils among skin leukocytes on Day 1 but contrary to the kinetics of pro-inflammatory cells, it remained significant for 3 weeks (Figure 1A, B). The peak of neutrophil accumulation can be used to define the onset of the resolution phase, which occurred between Day 1 and 7 in this model. These kinetics were confirmed by the quantification of pro-inflammatory cytokines (IL-6, TNFα) and monocytes attracting chemokines (CCL2, -3, -4) in the skin (Figure 1C). We did not detect any significant increase in interferons (IFNα and γ), usually associated with anti-viral or anti-bacterial Th1 responses, which supports that the wound healing response developed in the absence of any infection in this model. Conversely, IL-4 levels, associated with Th2 responses, increased during the resolution phase (Figure 1C). Th2 cytokines induce M2-like macrophages known to drive the resolution of inflammation and tissue repair. In concordance, the expression of genes related to M2-like macrophages (*Arg1*, *Retlna*), tissue repair macrophages (*Mgl2*), or the resolution (*Alox15*) tended to increase during the resolution phase (Supplementary Figure 1).

**Figure 1:**
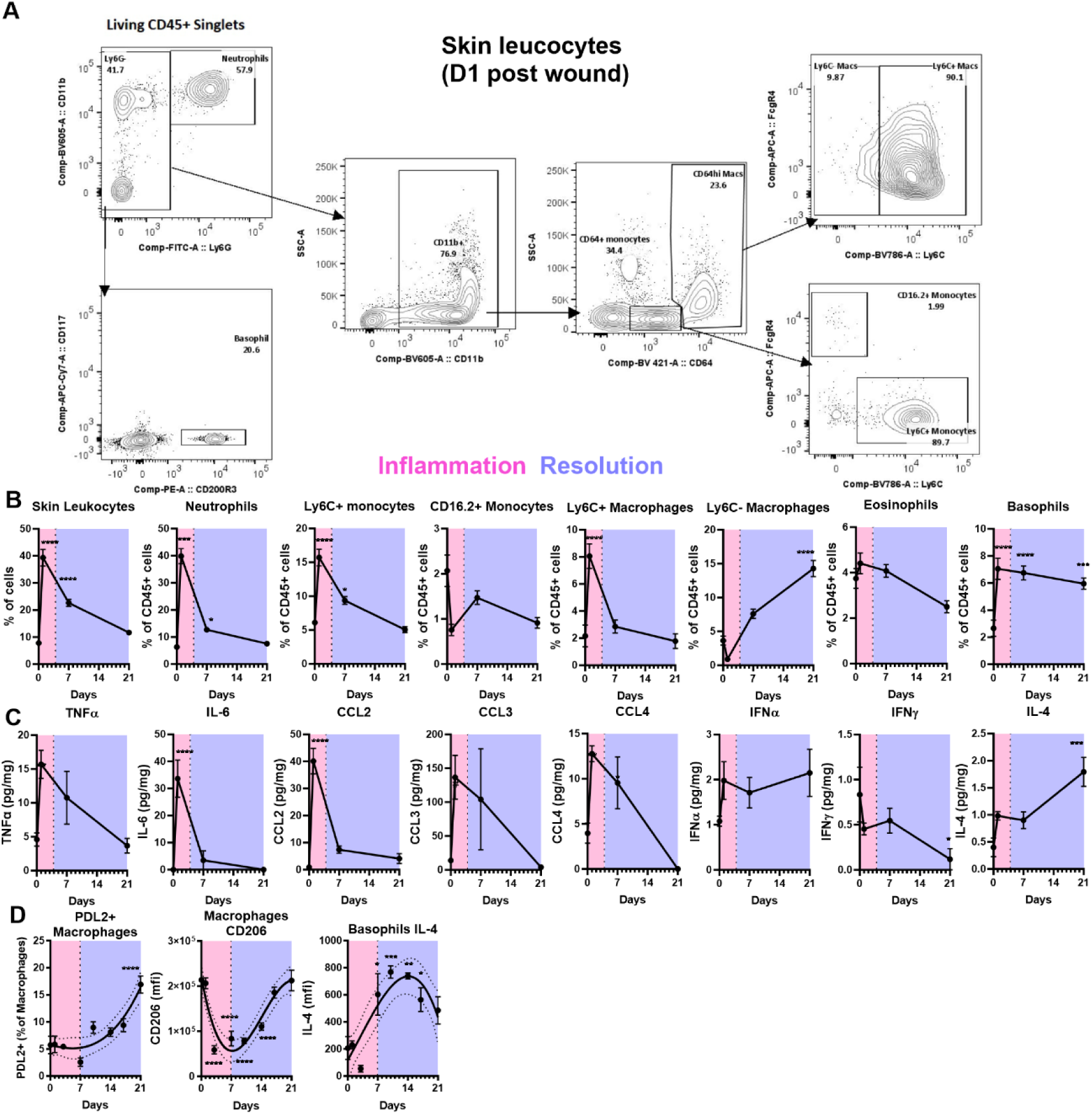
Basophils are enriched and activated during the resolution of skin wound healing. A-C) Wild-type mice were wounded by a sterile 2mm punch biopsy of the ear at day 0. A) Representative gating strategy of skin leukocytes among Living CD45+ Singlets and B) their proportions over time (n=23-40). C) Quantification of cytokines and chemokines by multiplex immunoassay over time, normalized by ear skin protein content (n=4-5). D) The phenotype of skin CD45+ CD64+ CD11b+ macrophages and CD45+ YFP+ basophils was analyzed by flow cytometry overtime after a 2mm plier-style ear punch in Basoph8x4C13R mice (n=4) as previously described(Ferrer-Font *et al*., 2020). A 3rd order polynomial interpolation with a 95% confidence interval is represented. Results are pooled from 3 to 10 independent experiments (B, C) or from a unique experiment (D). The inflammation and resolution phases are represented in red or blue with a day 7 cutoff. Statistics are ordinary one-way ANOVA corrected by a Holm-Sidak’s multiple comparison test against day 0. p<0.05: *; p<0.01: **; p<0.001: ***; p<0.0001: ****

We confirmed these results in a second model of sterile wound healing induced by a 2mm plier-style ear punch (used commonly for ear tagging), which induces a laceration but not a clean incision of the tissue (Rajnoch *et al*., 2003). This model showed similar yet delayed leukocyte infiltration kinetics which include an increased accumulation of basophils during the resolution phase (Supplementary Figure 2A). The resolution was characterized by an increase in M2-like macrophages expressing PDL2 and CD206 (Figure 1D). PDL2+ pro-resolution macrophages are known to differentiate from Ly6C+ inflammatory monocytes in response to basophil-derived IL-4 in the skin(Egawa *et al*., 2013; Miyake *et al*., 2024). Several immune cell types expressed IL-4 and/or IL-13 in the skin during the wound healing response, including mast cells, eosinophils, CD4+ T cells, and innate lymphoid cells, but their proportion or expression of type 2 cytokines did not evolve with inflammation or resolution kinetics (Supplementary Figure 2B). On the contrary, basophil IL-4 expression was strongly associated with the onset of the resolution phase (Figure 1D).

We then assessed by confocal microscopy where basophils locally infiltrated the skin during wound closure in the ear punch model (Figure 2A). We observed a strong presence of MCPT8+ basophils in the ear skin at D1 post-wound (Figure 2B). However, they were not accumulating as close to the wound edge as Ly6G+ neutrophils, or CD68+ monocytes/macrophages, but they rather settled at a certain distance instead, at a median of 670.1µm from the lesion (Figure 2B-D). We also confirmed the presence of numerous basophils in the ear skin by microscopy at D7 and D21 after injury (Figure 2E-F).

**Figure 2:**
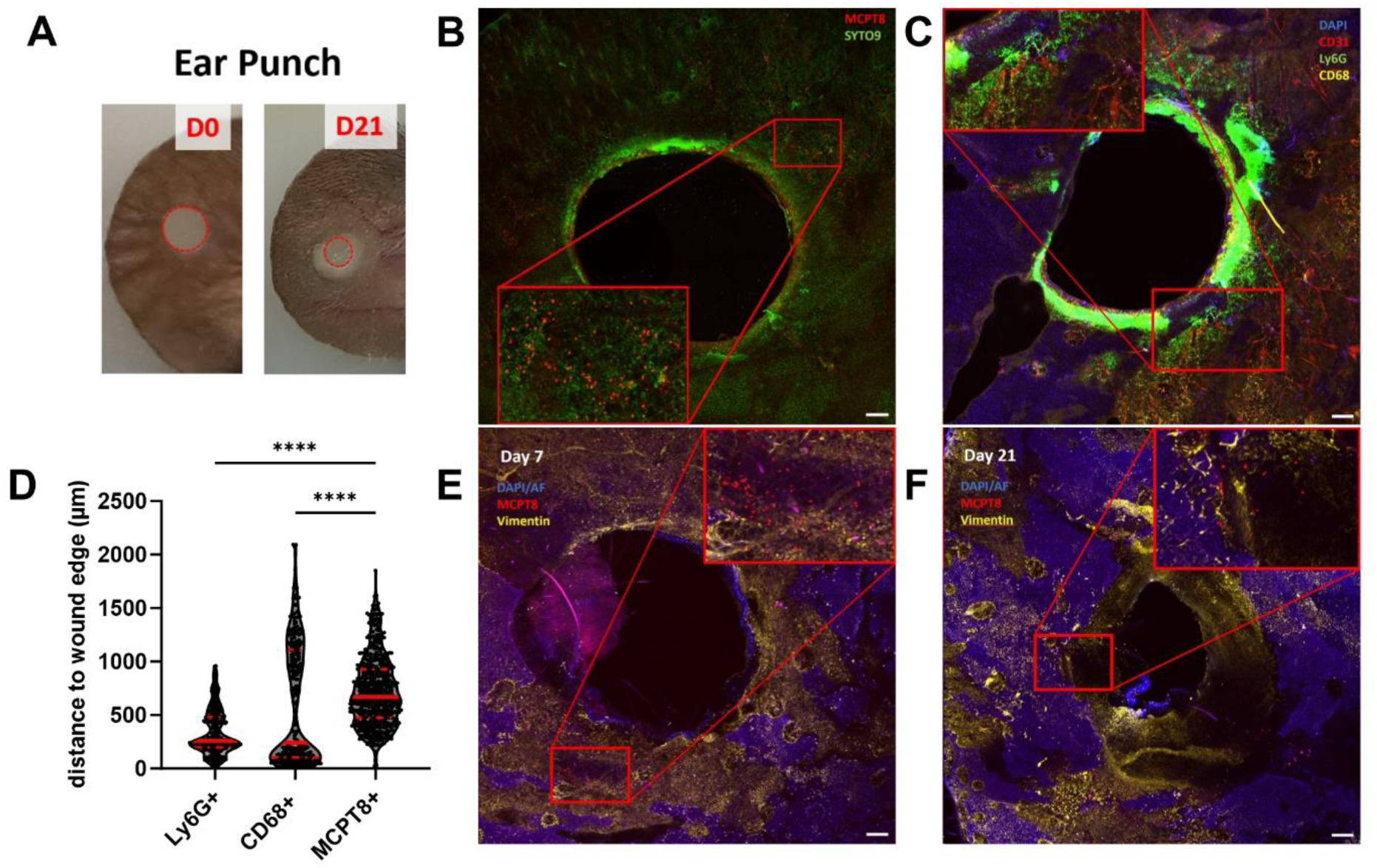
Basophils infiltrate ear skin wounds during inflammation and resolution. A) Representative photography of the area of the wound on day 1 and day 21 after ear punch biopsy (red dots). B-F) The ear skin was analyzed by whole mount confocal imaging of one dermal leaflet at (B-D) day 1, E) day 7, or F) day 21 after ear punch. B) The tdTomato fluorescence of a CTM8 mouse after counterstaining with SYTO9 without fixation or permeabilization and C) the expression of Ly6G, CD68, and CD31 of a wild-type mouse ear after gentle fixation and permeabilization and counterstaining with DAPI were used D) to quantify the closest distance of each of these cell types to the wound edge (n=360, 300, 559). E, F) The tdTomato fluorescence of CTM8 mice was used to identify basophils in the ear wounds at different time points under conditions of gentle fixation/permeabilization, alongside vimentin and DAPI counterstainings. B, C, E, F) Scale bars: 200µm. Each representation is a stitching of 9 images of the maximum projection of 3 to 5 overlapping z-stacks, at 10X. D) Results are pooled from three independent experiments. Statistics are a Kruskall-Wallis test corrected by a Dunn’s multiple comparison test against MCPT8+ cells. p<0.0001: ****

### 2) Basophils drive the resolution during the inflammation phase

Next, we studied the roles of basophils during the inflammatory phase of the wound healing response in the ear punch model by conditionally depleting them by injection of diphteria toxin (DT) in MCPT8-DTR mice. One day after wounding, basophil-specific depletion did not alter the skin immune infiltrate significantly (Figure 3A), except for an increased recruitment of CD16.2+ monocytes (Figure 3B). However, in the absence of basophils, monocytes, and macrophages exhibited a more activated phenotype: they expressed more CD11b (integrin and complement receptor), CD64 (activating high-affinity IgG receptor), and/or were enlarged (Figure 3B), and the skin microenvironment was more inflammatory and contained more pro-inflammatory cytokines and monocyte/macrophages attracting chemokines including IL-6, CCL2 and CCL4 (Figure 3C). As shown in Figure 1, there was a substantial decrease in both the leucocytic and the neutrophilic infiltrate one week after wounding, indicating the early resolution phase in this particular model. Basophil depletion from D-2 to D7 led to an increased accumulation of pro-inflammatory leukocytes in the wounds, including neutrophils and Ly6C+ macrophages without impacting the overall leucocytic infiltrate (Figure 3D). Moreover, neutrophils, Ly6C+ and CD16.2+ monocytes, and Ly6C+ macrophages showed increased activation markers and were all enlarged after basophil depletion (Figure 3E). Similarly, basophil depletion led to an accumulation of pro-inflammatory mediators at the onset of the resolution phase, including CCL2 and CXCL10 (Figure 3F). Overall, these results indicated that basophils dampen the activation of pro-inflammatory cells and wound inflammation during both the inflammation phase and the early resolution phase.

**Figure 3:**
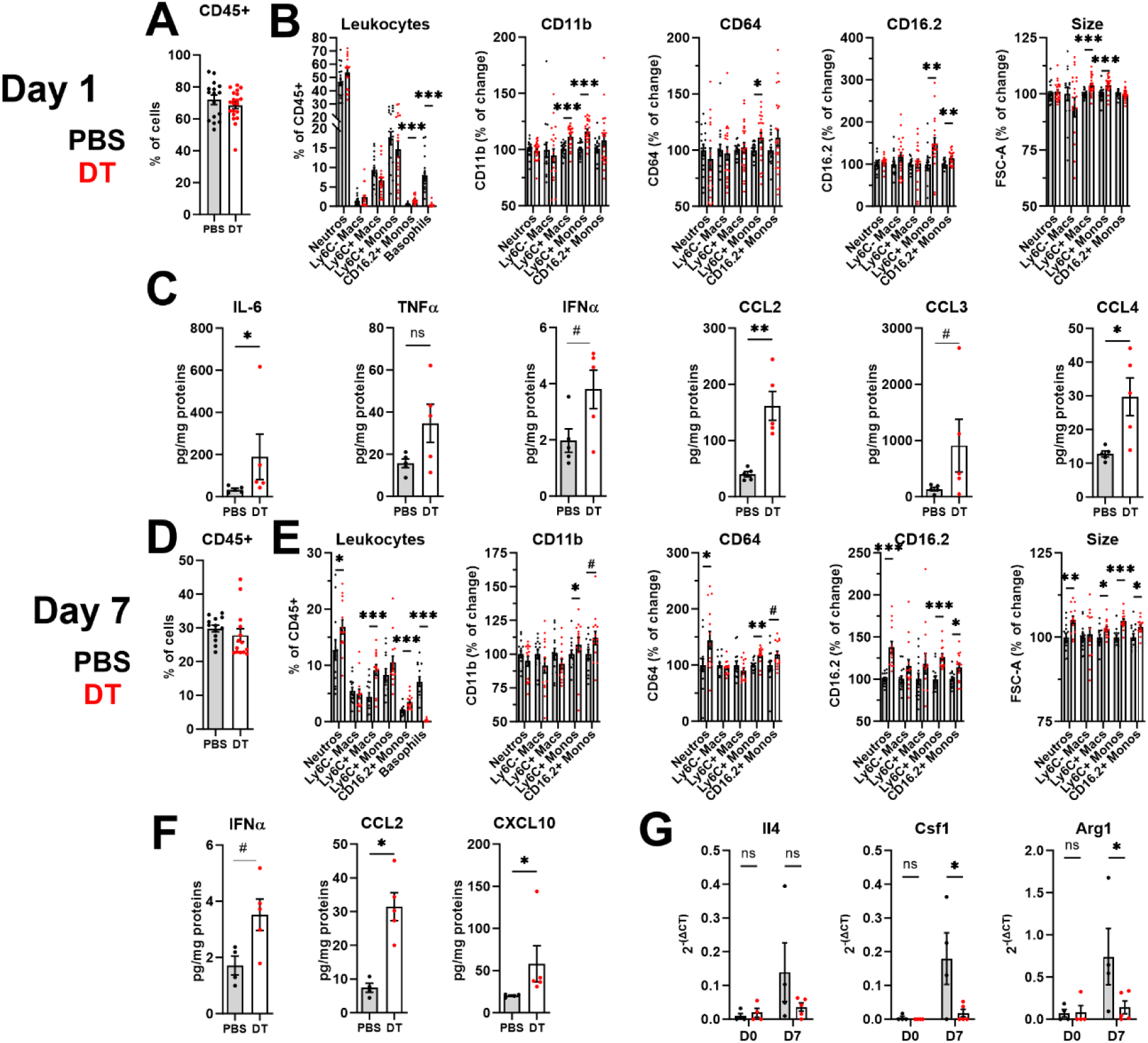
Basophils dampen activation of pro-inflammatory cells and inflammation during the first week of wound healing. MCPT8-DTR mice were depleted specifically of basophils by diphteria toxin injections A-C) at days -2, -1, and D-G) on day 5 post-wound. The infiltration of leukocytes, their proportions, and their phenotype were analyzed by flow cytometry A, B) at day 1 (n=16-24) or D, E) at day 7 (n=13-14). B, E) Each phenotype has been normalized on the control group (PBS) of each individual experiment for each population before pooling. Similarly, ear skin cytokine or chemokine content was quantified by multiplex immunoassay at C) day 1 or F) day 7 post wound, after normalization on total skin proteins (n=4-5). G) Skin gene expression was analyzed at day 0 and day 7 after normalization on GAPDH expression (n=3-5). Results are pooled from three independent experiments. Statistics are two-tailed Mann-Whitney tests. ns: non significant; p>0.1:#; p<0.01: #; p<0.05: *; p<0.01: **; p<0.001: ***; p<0.0001: ****

### 3) Basophils promote keratinocyte differentiation and the wound healing response

Uncontrolled inflammation has been associated with delayed wound healing response and the emergence of chronic wounds(Krzyszczyk *et al*., 2018). The ear biopsy punch model allows to quantify wound closure kinetics by monitoring the ear area that remained open. Of note, we observed different kinetics depending on the status (“Conventional or Specific Pathogen Free”) of the animal facility with an increased wound closure in conventional facilities (Figure 4A). As basophils promoted a resolution environment, we explored if basophils were affecting the kinetics of wound closure. Indeed, mice constitutively deficient in basophils showed delayed wound closure during the second week of healing (Figure 4B). Similarly, specifically depleting from D-2 to D7 after wounding significantly and transiently decreased wound closure in the first week (Figure 4C).

**Figure 4:**
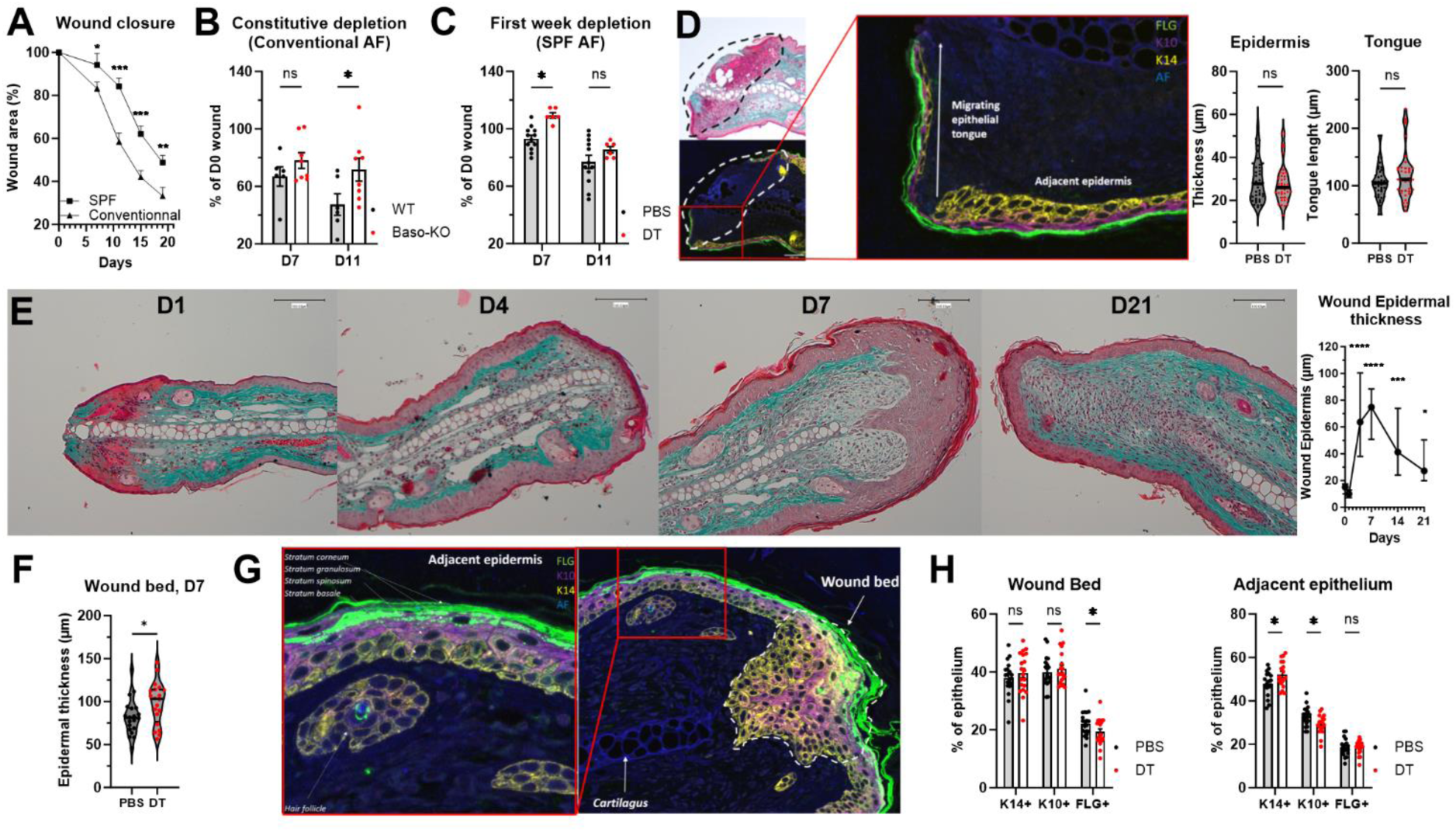
Basophils accelerate wound healing and the quality of the reepithelialization. Wild-type (WT) female mice with a C57BL/6J genetic background were bred in a specific pathogen-free (SPF, MCPT8^DTR^ mice) or conventional animal facility (AF, CTM8 mice), were ear punched at D0, and wound closure was monitored overtime until D19 (n=14-16). Similarly, wound closure at D7 and D11 was compared between B) CTM8 and CTM8xRosa-DTA (“Baso-KO”) mice bred in conventional AF (n=6-8) or C) MCPT8^DTR^ mice bred in SPF conditions depleted of basophils by injections of diphteria toxin (DT) at Day -2, -1, and Day 5 (n=6), or not depleted after injections of PBS (n=12). D) The thickness of the epidermis directly adjacent to the wound and the length of the migrating epithelial tongue were measured on Masson’s Trichrome colored ear sections at D1 post-wound in MCPT8^DTR^ mice treated with DT or PBS (n=36-40 wound edges, 2/ear). A D1 representative Masson’s trichrome coloration and a serial section stained with anti-filaggrin (FLG), Keratin 14 (K14), and Keratin 10 (K10) antibodies are depicted. AF (Blue) represents autofluorescence at 450nm. E) Similarly, the maximum wound bed epidermal thickness was measured over time and compared to D0 (n=16-32 wound edges, 2/ear) or F) measured at Day 7 in MCPT8^DTR^ mice treated with DT or PBS. G) Representative immunofluorescent staining of ear wound sections at Day 7, as in D, allowed the quantification of K14+ keratinocytes (stratum basale), K10+ keratinocytes (stratum spinosum), and FLG+ keratinocytes (stratum granulosum), as depicted. H) Analysis of the proportions of various keratinocytes in the wound bed and the adjacent epidermis at Day 7 as in G, in mice as in C). F, H) (n=20 wound edges, 2/ear). A, B, C, H) Means +/- sem or D, E, F) Medians +/- interquartile range are represented. Statistics are A) Repeated measure ANOVA or B, D) two-way ANOVAs with Holm-Sidak’s correction, or D, F) Mann Whitney tests or E) Kruskall-Wallis tests with Dunn’s correction or H) unpaired t-tests comparing the PBS to DT condition for each subset. ns: non significant; p<0.05: *; p<0.01: **; p<0.001: ***; p<0.0001: ****

As basophils are known to control keratinocyte differentiation during skin inflammation(Hayes *et al*., 2020; Pellefigues, Naidoo, *et al*., 2021; Strakosha *et al*., 2024), and the inflammatory processes in the wounds as shown here (Figure 3), we explored their roles in controlling reepithelialization in the ear punch model. Reepithelialization occurs via the proliferation, migration, and differentiation of keratinocytes, which will quickly cover the wound area by forming a “migrating epidermal tongue” from the adjacent epithelium which show increased proliferation and thickening. At 24h the wounds were never covered by keratinocytes in this model (Figure 4D). The thickness of the adjacent epidermis reflects the initial rate of keratinocyte differentiation and proliferation and the length of the migrating epithelial tongue can be measured as a surrogate for the rate of the reepithelialization(Nascimento-Filho *et al*., 2020; Bornes *et al*., 2021). The presence of basophils in the wounds at Day 1 was redundant for these phenomena (Figure 4D).

In this model, wounds were always covered by keratinocytes at D4, with a very heterogeneous epidermal thickness. The wound bed epidermis thickened heterogeneously until reaching a plateau at D7, before progressively returning to homeostasis levels (Figure 4E). Basophil depletion during the first week of healing led to increased inflammation and thickening of the wound bed epidermis at D7 (Figure 4F, G).

During homeostasis, Keratin 14+ keratinocytes (K14+) constantly proliferate in a “*stratum basale*” layer, before differentiating into K10+ keratinocytes (*Stratum spinosum*) and then expressing filaggrin (FLG+) in granules (*Stratum granulosum*), before dying by desiccation (S*tratum corneum)*. This controlled physiological differentiation of epidermal keratinocytes is altered during wound healing, with an increased proliferation and differentiation of keratinocytes in both the wound bed and the adjacent epidermis (Figure 4G). Non-healing chronic wounds show a defect in keratinocytes terminal differentiation and of their expression of both K10 and Flg, and they stay in a hyperproliferative state(Stojadinovic *et al*., 2008; Wikramanayake, Stojadinovic and Tomic-Canic, 2014). Here, one week post-wound, the epidermis of basophil-depleted mice was less differentiated: the wound bed contained fewer FLG+ cells, while the adjacent epithelium contained less K10+ but more K14+ keratinocytes (Figure 4H).

Thus, basophils accelerate skin wound closure and promote the differentiation of keratinocytes during the resolution phase of wound healing, but are redundant for the initial wound reepithelialization.

### 4) Basophils actively drive the resolution during the resolution phase

To decipher the role of basophils in controlling the immune response during the resolution phase of wound healing, we analyzed the skin infiltrate of mice constitutively deficient in basophils (“Baso-KO”), at 3 weeks post-wound (D21) in the ear punch model. While these mice did not show an aberrant quantity of leukocyte infiltration, basophil depletion led to a significant reorganization of the infiltrate, containing more pro-inflammatory cells, including neutrophils and Ly6C+ monocytes at the expense of non-inflammatory Ly6C-macrophages (Figure 5A). As this accumulation of pro-inflammatory cells during the resolution phase could be caused by the absence of basophils during the inflammatory phase (Figure 3), we used MCPT8-DTR mice to deplete basophils during only the 3^rd^ week of the wound healing response (Figure 5B). This conditional depletion, like the constitutive depletion, led to an accumulation of pro-inflammatory cells in the skin, including neutrophils and Ly6C+ macrophages, but not Ly6C+ monocytes (Figure 5C) or CD16.2+ monocytes (Supplementary Figure 3A). In addition, it led to the activation of wound neutrophils, Ly6C+ monocytes, and macrophages, which showed an increased expression of CD11b, CD16.2, CD64, and/or were enlarged. Ly6C-non-inflammatory macrophages did not show this activated phenotype after basophil depletion (Figure 5D). Importantly, this also led to an accumulation of CCL2, a pro-inflammatory chemokine involved in the recruitment and activation of Ly6C+ monocytes (Figure 5E). Unexpectedly, basophil conditional depletion did not lead to any decrease in the expression of M2-like markers at the surface of Ly6C-non-inflammatory macrophages (Figure 5F).

**Figure 5:**
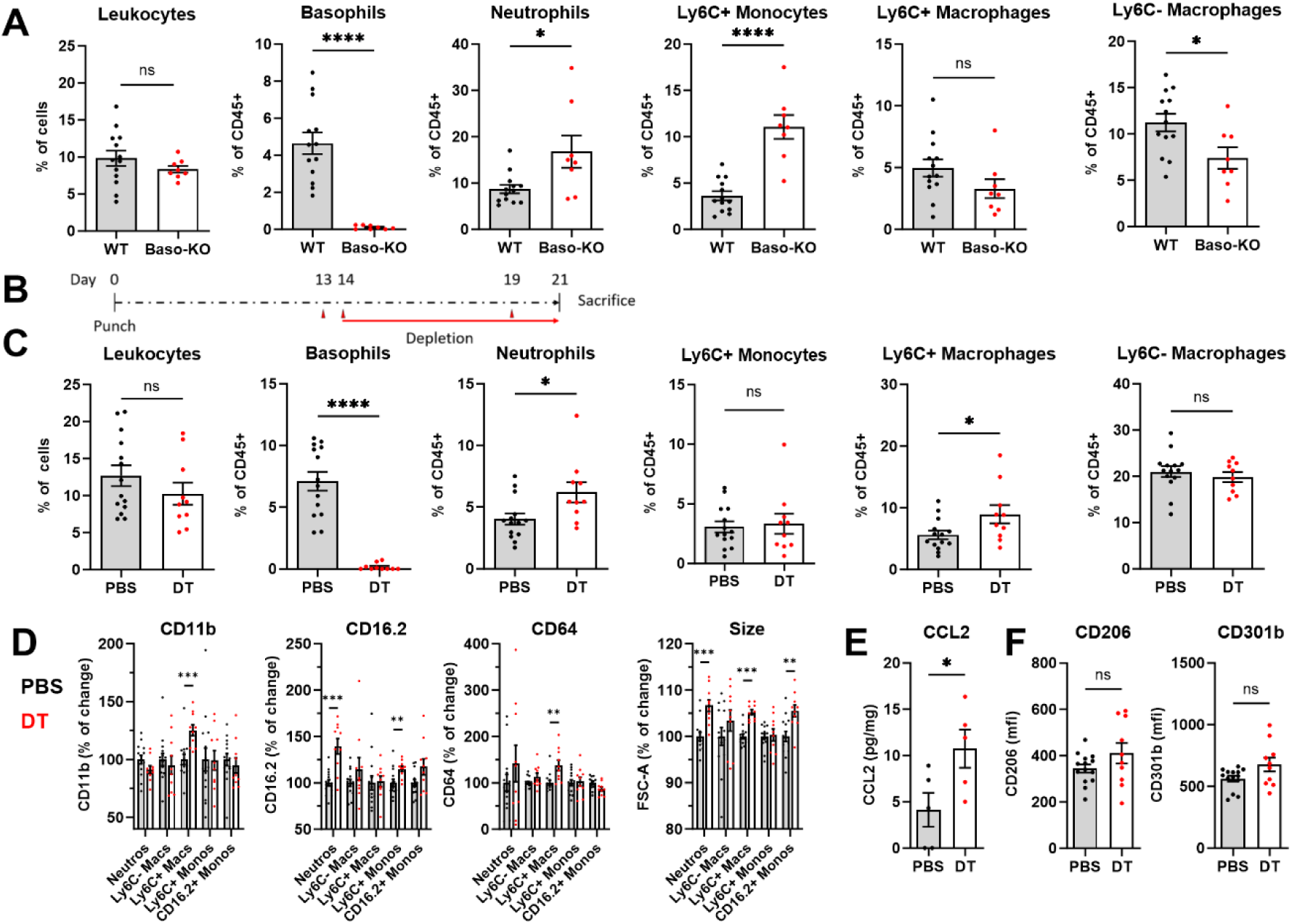
Basophils dampen the accumulation and activation of pro-inflammatory leukocytes during the resolution phase of skin wound healing. A) Wild type (WT) or basophil-deficient (Baso-KO) littermates were ear-punched and skin leukocytes were analyzed by flow cytometry at D21 (n=13, 8). B) Similarly, MCPT8^DTR^ mice were ear punched and injected with diphteria toxin (DT) or PBS on Days 13, 14, and 19 before being sacrificed on Day 21. C) The proportion of their skin leukocytes and D) their phenotype (normalized on control) were quantified by flow cytometry (n=14, 10). E) Similarly the chemokine CCL2 was quantified by immunoassay after normalization on total skin proteins (n=5) and F) the expression of CD206 and CD301b was quantified on Ly6C-macrophages by flow cytometry (n=14-10). Results are pooled from two to three independent experiments. Statistics are two-tailed Mann-Whitney tests. ns: non significant; p<0.05: *; p<0.01: **; p<0.001: ***; p<0.0001: ****

As basophils are recruited early during the inflammatory phase but remained in high proportions in the wounds during the resolution phase, we explored if basophils were actively recruited to the wounds during the resolution phase. We depleted basophils transiently during the resolution phase from D11 to D17 and then analyzed the skin immune infiltrate at D21 (Supplementary Figure 3B). Four days were sufficient for basophils to reinfiltrate the wounds to almost normal levels. No significant effect of their transient depletion on the accumulation of pro-inflammatory cells into skin wounds was observed (Supplementary Figure 3C). This suggests that basophils are actively recruited during both the inflammatory and the resolution phase into skin wounds to regulate transiently the immune environment.

Overall, these results show that basophils actively dampen inflammation during the late resolution phase of the skin wound healing response.

### 5) Basophil-derived IL-4 and CSF1 promote inflammation resolution and wound healing

As we previously showed that basophil-derived IL-4 and CSF1 were important for the resolution of skin allergic inflammation(Pellefigues, Naidoo, *et al*., 2021), we then explored if they were important for their pro-resolution properties during skin wound healing. We specifically deleted IL-4(Tchen *et al*., 2022) or CSF1(Pellefigues, Naidoo, *et al*., 2021) in basophils by Cre-Lox-specific recombination, and analyzed wound closure kinetics for 3 weeks. Here, basophil-specific depletion of CSF1 transiently delayed wound closure in the first week, while basophil-specific depletion of IL-4 delayed wound closure only during the third week (Figure 6A). This is in adequation with the fact that wound basophils only begin to express Il4 during the resolution phase (Figure 1D), while both blood and atopic skin basophils show a high constitutive expression of CSF1 (Uhlén *et al*., 2015; Pellefigues, Naidoo, *et al*., 2021).

**Figure 6:**
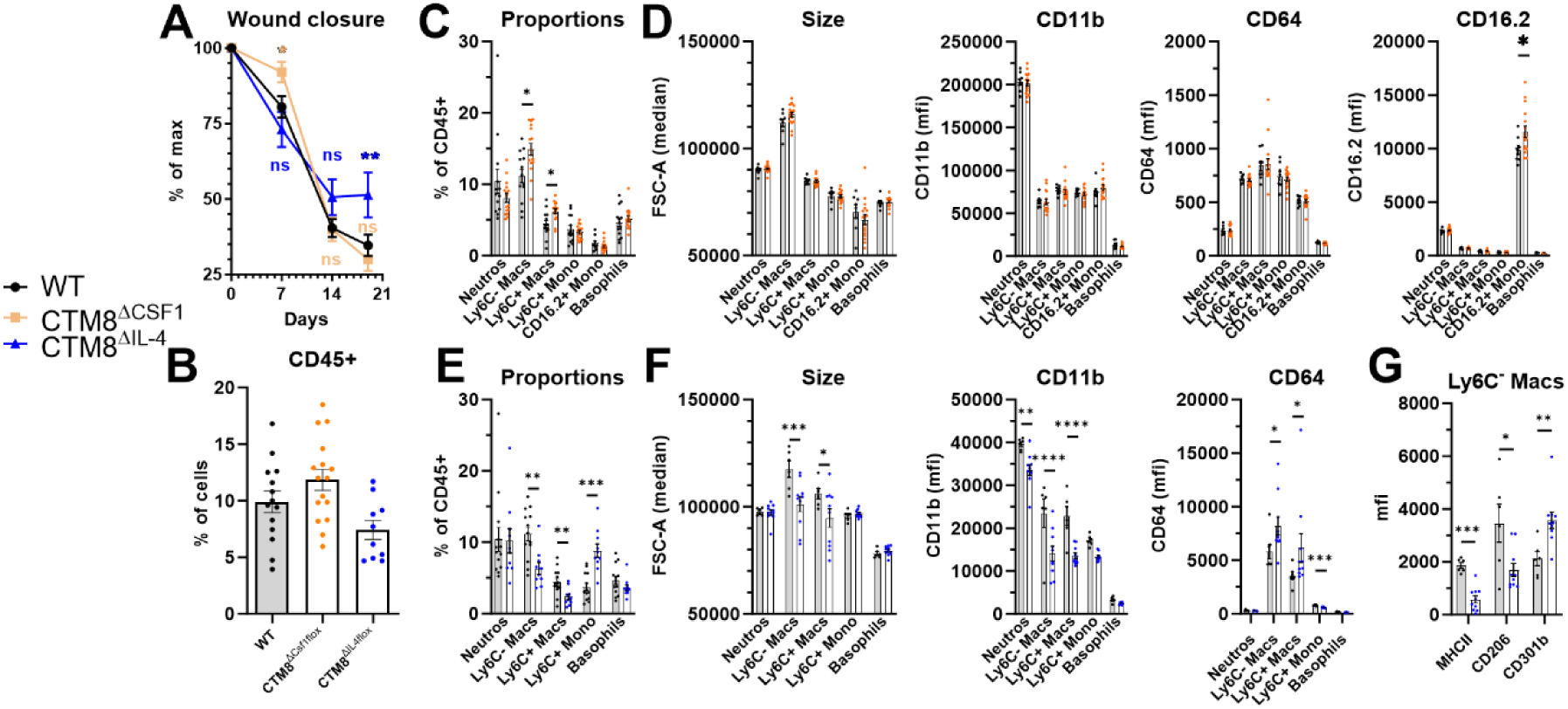
Basophils CSF1 and IL-4 independently promote wound closure and show complex immunoregulatory properties. Wild-type (WT) mice and mice with a basophil-specific deletion in Csf1 (CTM8^ΔCSF1^) or Il4 (CTM8^ΔIL4^) expression were ear-punched at Day 0. A) Ear hole area was monitored overtime after normalization on day 0 (n= 19, 21, 10), and B) total leukocyte content was determined by flow cytometry on Day 21 (n=14, 16, 10). The proportion and phenotype of skin leukocytes were analyzed by flow cytometry for C, D) CTM8^ΔCSF1^ or E, F, G) CTM8^ΔIL4^ mice. G) The expression of various surface markers by Ly6C-macrophages is represented. Neutros: Neutrophils. Macs: Macrophages. Mono: Monocytes. Results are A-C, E) pooled from 2 to 4 independent experiments or D, F, G) from a single experiment. Statistics are A) a two-way ANOVA with a Holm-Sidak’s post-test or C-G) Mann Whitney’s test to the WT condition. p<0.05: *; p<0.01: **; p<0.001: ***; p<0.0001: ****

As basophil-derived CSF1 deletion delayed wound closure at D7, we analyzed its immunoregulatory role at this time point. Deleting CSF1 in basophils had no significant impact on the overall infiltration of leukocytes (Supplementary Figure 4A), but was associated with a minor increase of CD16.2 expression by CD16.2+ monocytes (Supplementary Figure 4B).

We then analyzed the outcome of deleting CSF1 or IL-4 in basophils at three weeks post-wound, which had no significant impact on total leukocyte accumulation (Figure 6B). Deleting basophil CSF1 led to an accumulation of both Ly6C+ and Ly6C-macrophages in the skin at 3 weeks post-wound (Figure 6C). Furthermore, while it did not lead to any detectable change in Ly6C+ monocyte or macrophage activation, it increased the expression of the Fc receptor CD16.2 on CD16.2+ monocytes infiltrating the wounds (Figure 6D), as for Day 7 (Supplementary Figure 4B).

Deleting IL-4 in basophils promoted an accumulation of pro-inflammatory Ly6C+ monocytes in the wounds at D21, at the expanse of both Ly6C+ and Ly6C-macrophage populations (Figure 6E). Both macrophage subsets showed a smaller size and lower CD11b expression in the absence of basophils IL-4, similar to Ly6C+ monocytes. Furthermore, basophil-derived IL-4 deletion increased CD64 expression on both macrophage subsets but decreased it on Ly6C+ monocytes (Figure 6F). This was associated with a decrease of MHCII and CD206 expression, and an increase in CD301b expression, on Ly6C-macrophages, markers associated with M2-like polarization and wound healing (Figure 6G). This supports the known role of IL-4 on primary myeloid cells to downregulate CD64 and to upregulate CD11b and CD206 expression(Wong *et al*., 1992; de Waal Malefyt *et al*., 1993), and to promote macrophage differentiation from inflammatory monocytes(Finlay *et al*., 2023; Miyake *et al*., 2024).

Taken together, basophils show complex immunoregulatory properties and promote wound closure in part via their secretion of CSF1 and/or IL-4, albeit with different kinetics. Basophil secretion of IL-4 was particularly associated with a monocyte-to-macrophage transition and an M2-like polarization in the wounds.

### 6) Basophils are more activated and increased in skin wounds of aged mice

Aging is associated with a low-grade chronic systemic inflammation termed “Inflammaging”(Franceschi *et al*., 2018), and a dysfunction of the immune system involving in part defects in the resolution of inflammation. This defect in resolution is thought to be critical for the frequent development of chronic wounds observed in the elderly(De Maeyer *et al*., 2020). As basophils were scarcely studied in aged mice(van Beek *et al*., 2018), we explored the phenotype of blood leukocytes in 75 to 85 weeks old C57BL/6J mice (“Aged”). As expected, these mice tended to have fewer circulating leukocytes (Figure 7A) and showed decreased proportions of lymphocytes at the benefit of circulating monocytes and granulocytes populations, including neutrophils, Ly6C+ monocytes, CD16.2+ monocytes, and basophils (Figure 7B). Circulating basophils from aged mice displayed an increased activation status and notably showed increased IgE binding, without a detectable significant increase in the expression of the alpha chain of the high affinity receptor for IgE. Basophils in aged mice also showed an increased expression of CD11b and CXCR4 (chemokine receptor of CXCL12)(Pellefigues *et al*., 2018) and tended to express more CD200R1. They also displayed a decreased expression of CD200R3, which decreases upon basophil activation (Iwamoto *et al*., 2015; Pellefigues, Mehta, *et al*., 2021). This increased activation of basophils due to aging was also observed in the skin 24h after wounding (Figure 7D).

**Figure 7:**
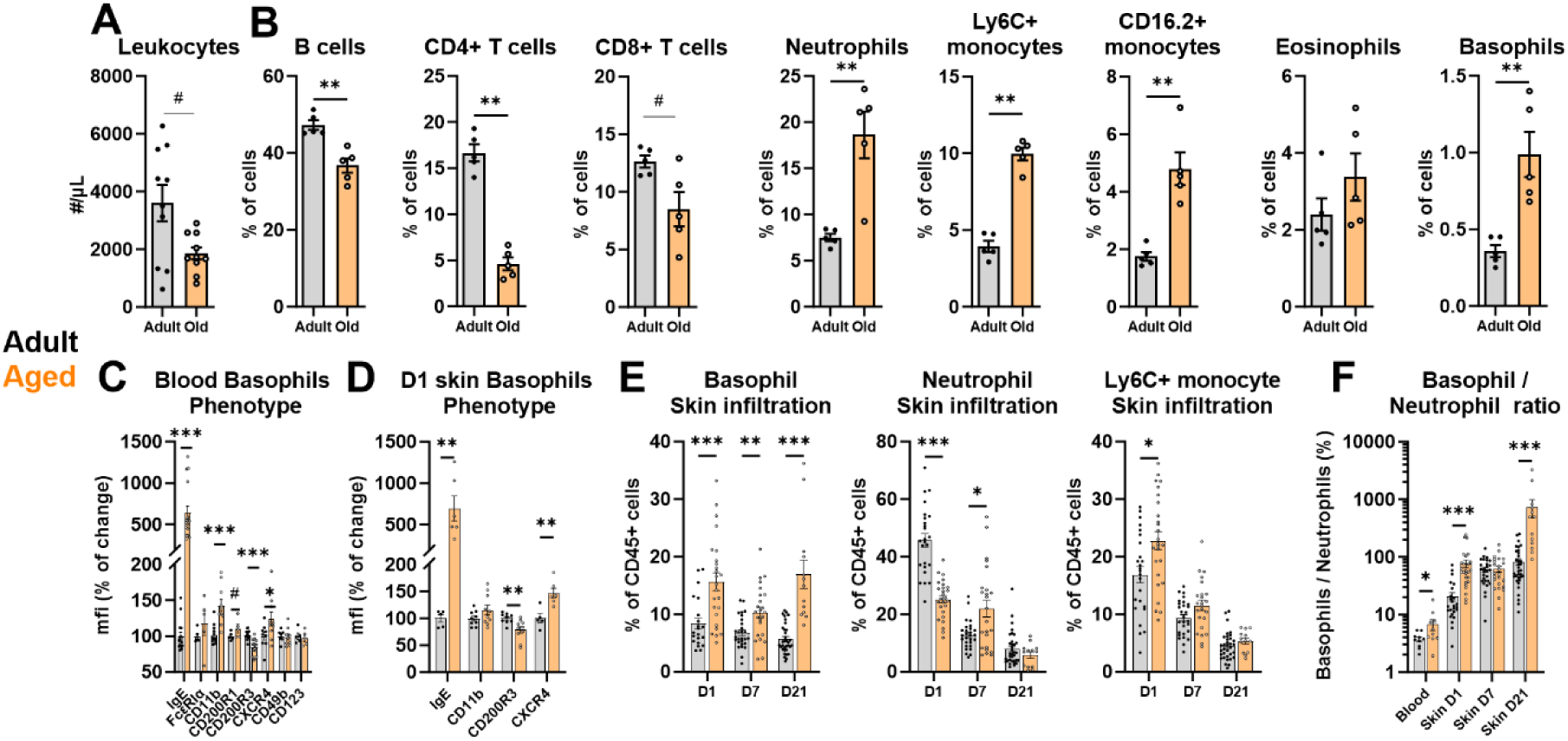
Basophils are more activated and infiltrate more skin wounds in aged mice. Naïve adult (8 to 14 weeks old) and Aged (75 to 90 weeks old) C57BL6/J mice were analyzed for their blood content in A) total leukocytes (n=10) and B) proportions off their main leukocyte subsets (n=5). Similarly, the surface phenotype of circulating basophils was analyzed in naïve mice (n=5-16), and D) in skin basophils 24h post wound (n=5-10). Similarly, E) basophil, neutrophil, and Ly6C+ monocyte skin infiltration were compared at different times after wounding (n=13-34) and F) a Basophil to Neutrophil ratio was calculated as a percentage for each condition (n=10-34). Results are from B) one single experiment or pooled from A) two to C-F) six independent experiments. Statistics are two-tailed Mann-Whitney tests to the adult condition. p<0.05: *; p<0.01: **; p<0.001: ***; p<0.0001: ****

After wounding basophils were more potently infiltrating the skin in aged mice, both during the inflammatory and the resolution phases (Figure 7E). In parallel, we detected a decreased infiltration of neutrophils while Ly6C+ monocytes were increased at 24h in aged mice. However, the proportion of neutrophils remained high at 7 days post-wound in aged mice (Figure 7E), supporting the known altered kinetics of neutrophil accumulation in the skin wounds of aged individuals(De Maeyer *et al*., 2020). As both basophil and neutrophil proportions were increased in the bloodstream of aged mice, we calculated the basophil-to-neutrophil ratio in skin wounds to appreciate the relative enrichment of basophils in the skin. This ratio showed a selective enrichment of basophils from the blood to the skin from day 1 to day 21 post-wound, and from adult to aged mice at days 1 and 21 (Figure 7F). Thus, aging is associated with an increased recruitment and activation of basophils during both the inflammation and resolution phases of wound healing.

Next, we sought to confirm these findings on another model of wound healing by reanalyzing our recently published scRNAseq dataset of the full-thickness dorsal wounds of adult (8 weeks old) and aged mice (88 weeks old) at days 0, 4, and 7(Vu *et al*., 2022). A t-SNE unsupervised clustering allowed to discriminate a cluster showing high *Kit* expression (“Mast cells”), from a second cluster expressing *Mcpt8,* a specific marker for basophils (Figure 8A). Both clusters were the main sources of *Il4* in the skin, while *Il13* and *Csf1* showed a broader pattern of expression. The basophil cluster also contained most of the cells positive for *Hgf*, a growth factor important for inflammation resolution and wound healing(Rutella, 2006; Nishikoba *et al*., 2020) and for *Alox15*, a lipoxygenase-producing specialized pro-resolution mediators (SPMs) and known marker of pro-resolution cells(Serhan, 2014). Basophils also expressed high levels of *Alox5ap*, an accessory protein fostering lipoxygenase activity, including SPMs production (Serhan, 2014) (Figure 8B). Analyzing the proportion of basophils in the wounds over time underlined that basophils infiltrate the wounds faster in aged mice (Figure 8C). As basophils are more activated early on in aged mice (Figure 7), we analyzed the kinetics of expressed genes related to resolution over time. At 4 days post-wound, basophils from aged mice tended to express more *Il4*, and expressed more *Csf1* than their adult counterpart, but showed a similar expression of *Hgf*. Importantly, the expression of *Ptgs2* (coding for COX2, an enzyme involved in lipid mediators and SPM production), and *Alox15* (whose expression is induced by type 2 cytokines(Serhan, 2014)), were mostly detected in wound basophils of aged mice (Figure 8D).

**Figure 8:**
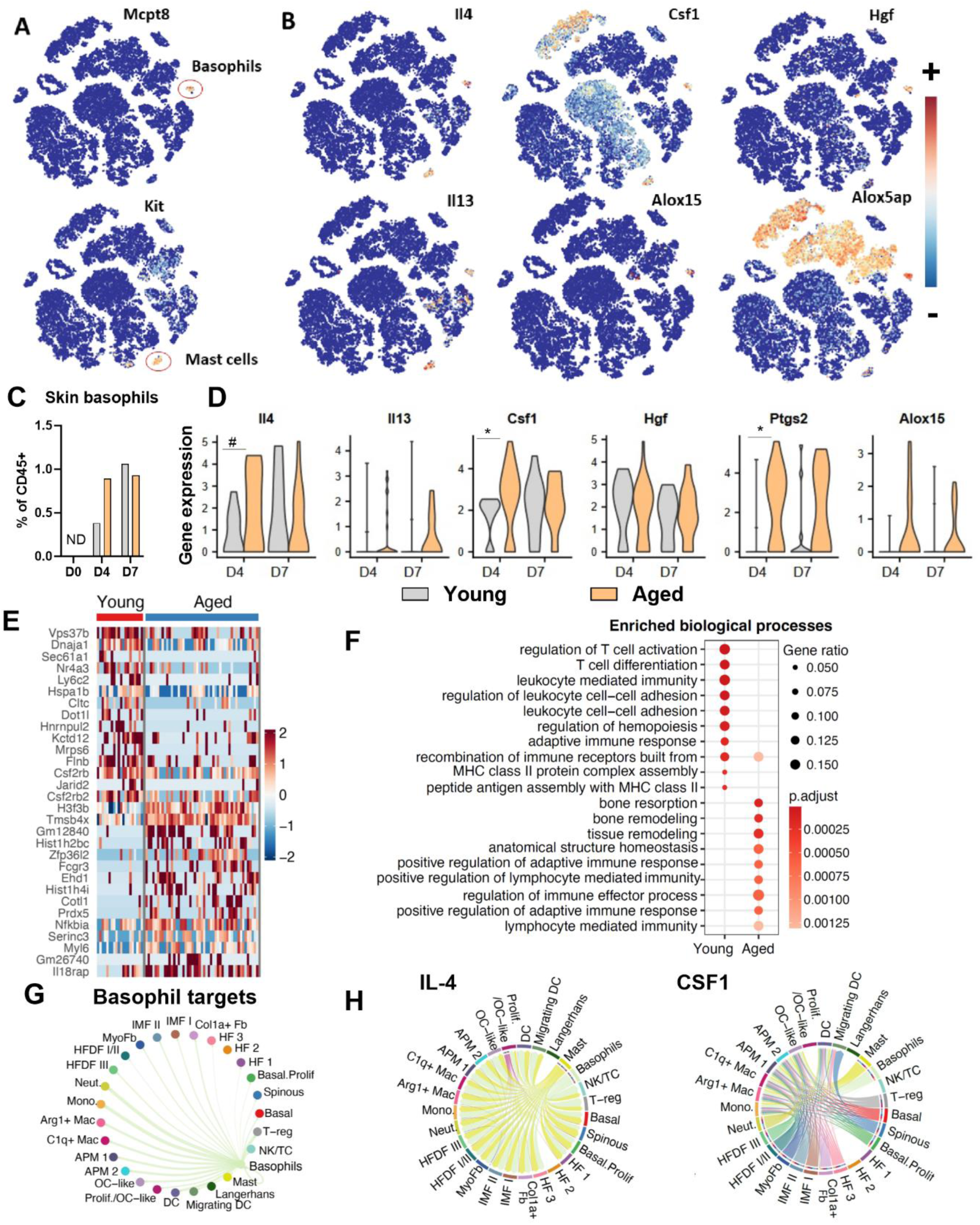
Basophils infiltrate dorsal wounds of aged mice and show a pro-resolution pro-remodeling signature. A previously published scRNAseq dataset (Vu *et al*., 2022) of dorsal wound cells from young (8 weeks old) and aged (88 weeks old) mice at 4 and 7 days post wounding (dpw) was reanalyzed for basophils. A) Representative expression of indicated genes after t-SNE unsupervised clustering identifies a “Basophil” cluster (MCPT8+ Kit-) from a “Mast cells” cluster (Kit+ MCPT8-) and B) their expression of relevant genes. C) The proportion of basophils was quantified in young and aged mice from the whole dataset. D) Basophil expression of selected genes from young or aged mice. E) Heatmap of the most differentially expressed genes between young and aged mice basophils (4dpw + 7dpw). F) Enriched gene ontology (GO) of biological processes between young and aged basophils at 4dpw. G) Circle plots of CellChat-predicted cellular targets of basophils (young + aged), and H) chord plots showing the inferred IL-4 or CSF1 signaling networks and cell types involved, in aged mice at 4dpw. Arrow width represents the predicted strength of the interactions. A, B) v2 + v3 datasets and C-G) v3 dataset only. D) Statistics are Mann-Whitney tests. #:p<0.1; *: p<0.05; ND: Not Detected. IMF: Immune modulating fibroblasts. MyoFb: Myofibroblasts. Fb: Fibroblasts. HFDF: Hair Follicle (HF)-associated dermal fibroblasts. HF1-3, Basal prolif., Basal and Spinous are keratinocytes subsets. OC-like: Osteoclast like. DC: Dendritic cell. APM: Antigen presenting macrophage. NK/TC: Natural Killer and T cell cluster.

The early activation of basophils from aged mice could be due to an intrinsic property related to age. To test this hypothesis, we analyzed the main differential gene expression between wound basophils in adult and aged mice. Adult basophils expressed more genes related to endocytosis (*Vps37b, Cltc, Flnb*), cellular stress or inflammation (*Dnaja1, Hspa1b, Hnrnpul2*), regulation of transcription or traduction (*Sec61a1, Nr4a3, Jarid2*), and the receptor for common β chain cytokines (*Csf2rb, Csf2rb2*), which include IL-3, a growth factor and cytokine critical for basophils homeostasis. Interestingly, they also show an increased expression of *Ly6C2*, which we did not observe by flow cytometry at the protein level in the ear punch model (Supplementary Figure 1A), but that can be induced in response to IL-3 *in vitro*. It was also observed in the skin in a model of atopic dermatitis(Pellefigues *et al*., 2019). By contrast, basophils from aged mice were characterized by an increased expression of histone variants (*H3f3b, Hist1h2bc, Hist1h4i*), known to increase during cellular senescence(Dubey, Dubey and Kleinman, 2024) as well as genes coding proteins with anti-inflammatory properties such as thymosin β4 (*Tmsb4x*), peroxiredoxin 5 (*Prdx5*) (Philp and Kleinman, 2010; Knoops *et al*., 2011) or IκBα (*Nfkbia*) (Pellefigues, Mehta, *et al*., 2021). They also expressed more coactosin-like protein (*Ctl1*), a chaperone stabilizing lipoxygenase activity (Basavarajappa *et al*., 2014), CD16 (*Fcgr3*), known to be expressed by a minor subset of basophils in human blood(Vivanco Gonzalez *et al*., 2020), and *Il18rap*, coding for a chain of the IL-18 receptor. IL-18 is an epidermal-derived alarmin known to potently activate basophils (Ferrucci *et al*., 2005; Pellefigues, Mehta, *et al*., 2021), which is increased in the plasma of both old humans and mice(Brigger *et al*., 2020) (Figure 8E). We then focused on the differences between adult and aged wound basophils at 4 days post-wound, which showed increased activation in aged basophils. Gene ontology (GO) of biological processes revealed a bias towards T cell activation and regulation of leukocyte adhesion in adult mice, whereas basophils from aged mice were enriched for genes associated with remodeling and homeostasis (Figure 8F). Inference of putative cell-cell communications based on ligand and receptor gene expression revealed that basophils’ primary targets were more likely to be neutrophils, monocytes, and macrophage subsets than other cell types in the wounds (Figure 8G). Furthermore, analyzing the intercellular signaling networks of IL-4 and CSF1 revealed basophils and mast cells to be the predominant sources of IL-4 in aged mouse skin wounds, able to interact with multiple different cellular targets, whereas CSF1 expression was broader, but able to target mainly cells of the monocytic lineage, including monocytes, macrophages subsets, osteoclast-like cells, and dendritic cells (Figure 8H).

These results confirm that basophils infiltrate mouse skin wounds and display a unique immunoregulatory potential during the resolution of inflammation. Overall, while basophils from aged mice display some markers of senescence, they show a faster activation and ability to infiltrate the wounds. Their transcriptomic signature is biased toward tissue remodeling and homeostasis when compared to adult basophils in the wounds. This suggests that basophils’ pro-resolution and pro-healing properties are not diminished but enhanced by aging.

### 7) Basophils drive the resolution in wounds of aged mice

We then analyzed the function of basophils during the inflammation, early resolution, and late resolution phase of wound healing in aged mice (75 to 85 weeks old) using the ear punch model. Depleting specifically basophils in aged mice did not change the overall leukocyte infiltrate at day 1 (Figure 9A). However, it led to a dramatic change in the quantity of the infiltrate, with a strong rise in neutrophilic infiltration at the detriment of Ly6C+ monocytes (Figure 9B). At this early time point, basophils from aged mice were dampening the activation of neutrophils, monocytes, and macrophages into the wound, as revealed by their increased activation phenotype (CD11b, Size, CD64, CD16.2) after basophil depletion (Figure 9C). Similarly, while basophil depletion during the first week did not change the quantity of the leukocyte infiltrate (Figure 9D), it promoted the accumulation of monocytes (Ly6C+ and CD16.2+), Ly6C+ pro-inflammatory macrophages (Figure 9E) and their activation alongside neutrophils (Figure 9F). Importantly, basophil depletion starting at D-2 led to a significant delay in ear wound closure at D7 (Figure 9G), indicating that basophils were accelerating wound healing in aged mice as well.

**Figure 9:**
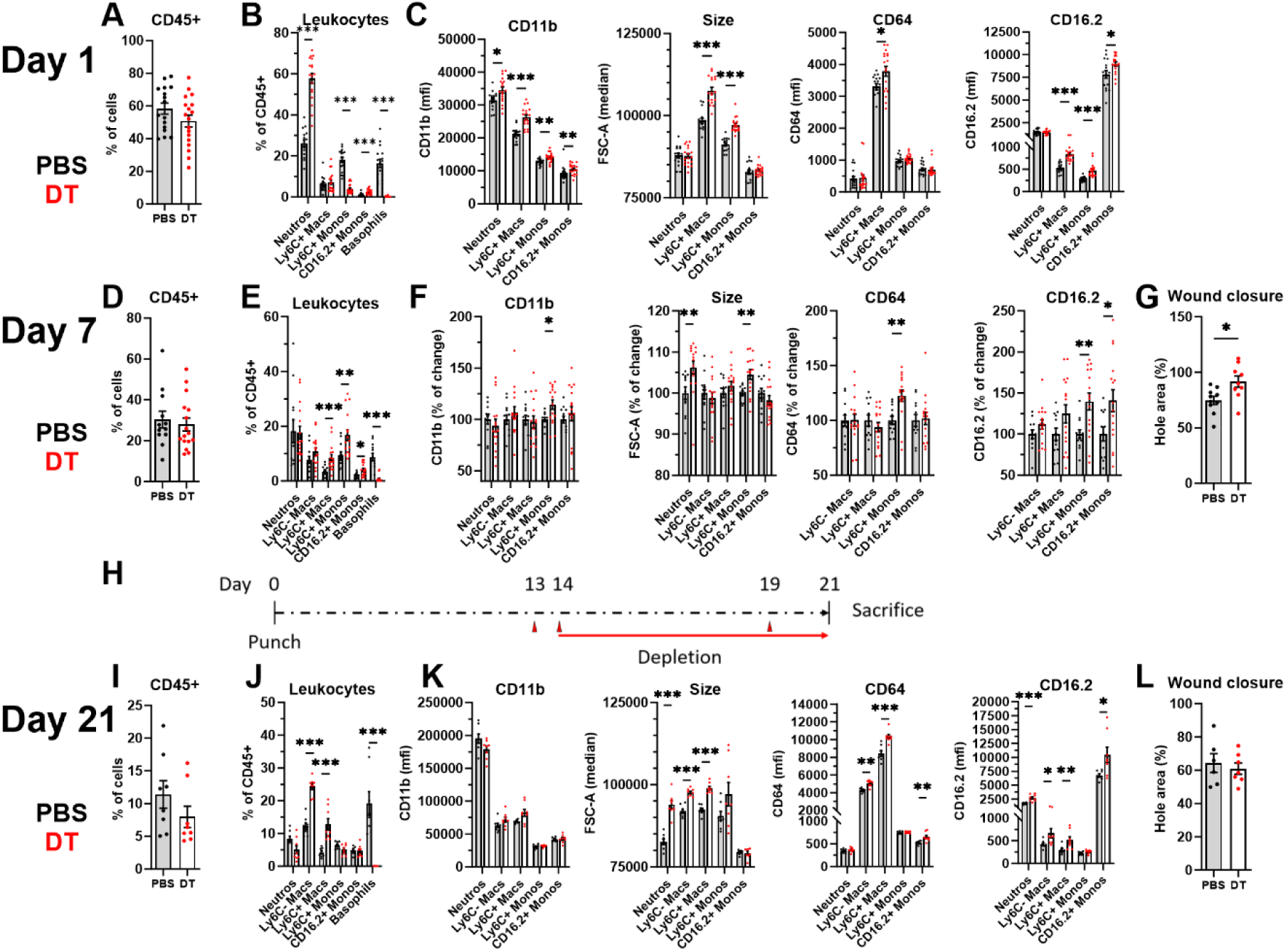
Basophils promote the resolution and wound closure in aged mice. Aged MCPT8^DTR^ mice were ear punched at Day 0 and depleted in basophils by injection of diphteria toxin at A-C) Day -2, -1, and D-G) Day 5, or H-L) at Days 13, 14 and 19 (as represented in H). Total skin leukocyte A, D, I) content and B, E, J) composition was analyzed by flow cytometry at Days 1, 7, or 21 as indicated, and the C, F, K) phenotype of selected cell types was analyzed. Wound closure was analyzed G) at day 7 in female mice (n=10, 9) and L) at day 19 in male mice (n=6, 7). Results come from A, B, C, I, J, K) one single experiment or D, E, F) are pooled from two independent experiments. F) Results have been normalized on each control condition. A-C) n=16, 18. D, F) n=12-16. I-K) n=8, 8. Statistics are two-tailed Mann-Whitney tests. p<0.05: *; p<0.01: **; p<0.001: ***; p<0.0001: ****

Next, we analyzed the role of basophils during the third week of the wound healing response in aged mice after their conditional specific depletion (Figure 9H). Basophil depletion did not lead to a significant decrease in skin leukocyte content (Figure 9I) but promoted the accumulation of both Ly6C- and Ly6C+ macrophages in the wounds (Figure 9J). The depletion further led to an activation of the neutrophil and monocyte/macrophage compartments (Figure 9K) but did not induce an observable delay in wound closure. Of note, this last analysis was limited to aged males (Figure 9L). Importantly, we observed that wound closure was faster in aged versus adult female mice, as previously shown(Nishiguchi *et al*., 2018), but also in aged females when compared to aged males (Supplementary Figure 5A).

Importantly, transiently depleting basophils from D10 to D17 in aged mice did not significantly decrease wound basophil numbers at D21, as in adult mice (Supplementary Figure 3D) suggesting that basophils quickly reinfiltrated the wounds during the resolution phase. This transient depletion led to a lasting accumulation of pro-inflammatory Ly6C+ macrophages and tended to increase the accumulation of neutrophils and Ly6C+ monocytes, in the wounds (Supplementary Figures 5B-D).

Basophils promoted an early neutrophil-to-monocyte transition in aged mice wounds before reducing monocyte and macrophage accumulation and activation for at least 3 weeks. This was associated with an acceleration of wound closure during the first week. Overall, basophils show pro-resolution and pro-healing properties in old mice wounds.

## Discussion

Basophils are pro-inflammatory cells involved in allergic diseases and protection against helminths, type 2 immune responses involving IgE reactivity (Miyake, Ito and Karasuyama, 2022; Poto *et al*., 2023). However, type 2 immune responses have also evolved as a response to tissue damage which regulates tissue remodeling and healing(Gieseck, Wilson and Wynn, 2018), and are well known to antagonize and counterbalance type 1 immune responses against pathogens, and their toxicity for the tissues(Gause, Wynn and Allen, 2013). Basophils are important players in this network. Besides antibody-dependent responses, they are also potently activated by innate immune mechanisms. They are particularly activated by epidermal-derived alarmins, which trigger the production of type 2 cytokines(Pellefigues, Mehta, *et al*., 2021), suggesting that they can participate in the response to tissue damage. Here, we demonstrate that basophils are strongly recruited to skin lesions in the absence of any noticeable infection. They display a particular wound infiltration kinetics: while pro-inflammatory leukocytes such as neutrophils infiltrate transiently the wounds during the inflammation phase, basophils accumulate in the wounds and persist through the resolution phase, as already shown in a model of atopic skin inflammation(Pellefigues, Naidoo, *et al*., 2021). The fact that a basophil-to-neutrophil ratio is higher in the wounds than in the blood demonstrates the relative specificity of basophil infiltration in the wounds during the first three weeks of the healing process. We also show that basophils infiltrate the skin farther away from the wound edge than pro-inflammatory cells. This particular localization suggests their pro-resolution function may help to contain the spread of inflammation close to the wound site. In line with a pro-resolution role, basophil depletion led to an increased inflammation characterized by the accumulation of the chemokine CCL2 and pro-inflammatory leukocytes expressing its receptor CCR2 (including neutrophils and classical monocytes), during both the inflammation and resolution phases. In agreement, basophils were previously shown to inhibit neutrophil recruitment during skin infection(Leyva-Castillo *et al*., 2021), epidermal abrasion(Strakosha *et al*., 2024), or atopic-like skin inflammation and resolution(Pellefigues, Naidoo, *et al*., 2021). Thus, it is tempting to speculate that basophil accumulation participates in shaping chemokine gradients around the wounds, thereby fine-tuning the recruitment of pro-inflammatory leukocytes at the inflammatory site and limiting unnecessary damage to the surrounding tissue.

Our data suggests that basophils accelerate ear wound closure between one and two weeks post-wound. Several biological processes participate in the regulation of the wound healing response, including hemostasis, re-epithelialization, angiogenesis, and tissue remodeling. As basophils are potently activated by epithelial-derived signals(Pellefigues, Mehta, *et al*., 2021), and regulate keratinocyte homeostasis in various models(Hayes *et al*., 2020; Leyva-Castillo *et al*., 2021; Pellefigues, Naidoo, *et al*., 2021; Strakosha *et al*., 2024), we focused on their role in the reepithelialization of skin wounds. Basophils did not control the initial phases of reepithelialization, known to be driven by IL-17A inflammation(Konieczny *et al*., 2022). Instead, we demonstrate that basophils are accelerating keratinocyte terminal differentiation in the wound bed during the resolution phase, notably at the time at which they promote wound closure. This is in concordance with recent observations that basophils promote keratinocyte differentiation, including their expression of the genes *Krt10* and *Flg*, by counteracting IL-17A-driven inflammation during the recovery phase of experimental epidermal inflammation(Strakosha *et al*., 2024). We previously described the role of basophils in promoting keratinocyte homeostasis during the resolution phase of atopic dermatitis-like inflammation (Pellefigues, Naidoo, *et al*., 2021). The differentiation of keratinocytes is critical for the quality of the wound healing response, as it restores the barrier function of the epidermis, which reduces microbial-driven local inflammation. Indeed, chronic wounds show hyperproliferative keratinocytes lacking the expression of terminal differentiation genes such as *Flg* and *Krt10*(Stojadinovic *et al*., 2008; Wikramanayake, Stojadinovic and Tomic-Canic, 2014). Further studies will be necessary to determine if the control of keratinocyte homeostasis by basophils in skin wounds involves their direct secretion of IL-4, as suggested in a model of *S. aureus* infection(Strakosha *et al*., 2024), or indirect mechanisms such as the control of the local inflammatory milieu. In agreement with the latter, we did not observe basophils close to the epidermis, and wound basophils seem biased to interact directly more with immune cells from the granulocyte–monocyte lineage than with keratinocytes (Figure 8G).

CSF1 is a critical regulator of monocyte/macrophage homeostasis through its interaction with the CSF1R. CSF1 is expressed constitutively at high levels by basophils (Uhlén *et al*., 2015; Pellefigues, Naidoo, *et al*., 2021) and many types of mesenchymal cells(Sehgal, Irvine and Hume, 2021). In the skin, CSF1 can promote monocyte/macrophage expansion and foster an M2-like bias in some particular models(Braza *et al*., 2018; Pellefigues, Naidoo, *et al*., 2021), as well as the resolution of inflammation during wound healing(Klinkert *et al*., 2017; Kapanadze *et al*., 2023). Similarly in our ear punch model, basophil-derived CSF1 accelerated wound closure during the first week and dampened the accumulation of Ly6C+ macrophages at 3 weeks post-wound. In the dorsal wound model, CSF1 expression was increased in basophils from aged mice at day 4 post-wound. Overall, basophil-derived CSF1 seemed to influence the differentiation of wound macrophages, which was important to accelerate wound closure during the first week of healing. The specific deletion of basophils’ IL-4 induced an accumulation of Ly6C+ monocytes and a decrease of both subsets of macrophages analyzed decreasing at the same time their expression of some M2-like markers. IL-4 promotes macrophage proliferation *in situ*(Jenkins *et al*., 2011, 2013) and a monocyte-to-macrophage phenotypic transition(Wong *et al*., 1992; de Waal Malefyt *et al*., 1993; Dang *et al*., 2023; Finlay *et al*., 2023). While testing these hypotheses was out of the scope of this study, we could identify a key role for basophil-derived IL-4 to promote macrophage homeostasis, to cause an M2-like bias during the resolution phase, and to accelerate wound closure at three weeks post-wound. IL-4 signaling is critical to limit age-induced inflammation at the organism and macrophage levels(Franceschi *et al*., 2018; Zhou *et al*., 2024), and notably promotes a central immunosuppressive myelopoiesis program in a model of chronic inflammation(LaMarche *et al*., 2024). In this light, basophil’s IL-4 secretion may be a key natural regulator counterbalancing monocyte/macrophage-dependent chronic inflammation. Future studies will be needed to decipher how basophil-derived IL-4 acts locally at the inflammatory site, or systemically by influencing myelopoiesis.

The phenotype of basophils from aged mice has only been scarcely studied. One seminal study found increased numbers of basophils in the spleens and bone marrow of C57BL/6 aged mice(van Beek *et al*., 2018). We confirm the peripheral basophilia in aged mice and that aged mice basophils exhibit a decreased expression of CD200R3, as it is observed upon basophil activation. In addition, we found aged mice basophils express more CD11b, CXCR4, and IgE in the blood, while FcεRIα, the high-affinity IgE receptor remains stable. This last finding seems counterintuitive as most FcεRIα receptors are thought to be occupied by IgE *in vivo*, and as IgE binding stabilizes the membrane expression of FcεRIα (Turner and Kinet, 1999). However, previous studies show that germ-free or antibiotics-treated miceshow high serum IgE, peripheral basophilia, and increased basophil IgE binding without any noticeable increase in FcεRIα expression(Hill *et al*., 2012). As age-associated microbiota can alter the phenotype of basophils(van Beek *et al*., 2018), it may have increased the binding of IgE to basophils in a similar manner. Importantly, basophils showed potent anti-inflammatory, pro-resolving, and pro-healing properties in aged mice, while displaying a transcriptomic signature biased towards tissue remodeling and homeostasis (Figures 8-9). This is unexpected, as aging induces a chronic low-grade inflammation state arising from cellular senescence and a defect in the resolution(Franceschi *et al*., 2018). During wound healing, macrophages fail to adopt a pro-resolutive state and to perform efferocytosis, which facilitates the accumulation of neutrophils and the establishment of non-healing chronic wounds(De Maeyer *et al*., 2020; Dube *et al*., 2022). Aging is also characterized by a defect in type 2 cytokine signaling, while exogenous IL-4 restores age-induced macrophage defects, and a healthy lifespan, in mice(Zhou *et al*., 2024). Thus, basophil secretion of IL-4 in aged mice appears as a mechanism arising to counterbalance age-associated inflammation. The fact that basophils express ALOX15 in the wounds of aged mice suggests basophils may further promote the resolution of inflammation through the production of SPMs(Basil and Levy, 2016). Old age is characterized by an increasing interindividual variability, and old mice exhibit a more variable wound healing rate than adult mice(Mahmoudi *et al*., 2019). Our observations suggest that this variability may be due partly to basophil infiltration and that individuals defective in basophil pro-resolving properties would be more susceptible to the establishment of chronic wounds.

Overall, we identify basophils as pro-resolving cells regulating wound inflammation, particularly in the context of aging. Harnessing these properties may lead to new therapeutic strategies to prevent chronic wounds development in the elderly.

## Methods

### Mice and treatments

MCPT8^DTR,^(Wada *et al*., 2010) , CTM8(Tchen *et al*., 2022), IL-4^LoxP^ and CSF1^Loxp^ mice(Shibata *et al*., 2018; Pellefigues, Naidoo, *et al*., 2021) were bred and maintained on a C57BL/6J background. C57BL/6J and Rosa-DTA mice were from The Jackson Laboratory through Charles River Laboratories or Envigo. “Baso-KO” mice are CTM8xRosa-DTA(Tchen *et al*., 2022). Littermates were used to compare mice of the same sex and age in individual experiments. For experiments related to wound closure kinetics, we found sex-related significant differences and thus analyzed male and female kinetics independently. Sex bias is always described in the figure’s legend. MCPT8^DTR^ mice were bred and maintained in specific pathogen-free conditions, or conventional conditions for experiments involving Cre-Loxp recombination, as described in the figure’s legend. Most of the study was conducted in accordance with the French and European guidelines and approved by the local ethics committee “Comité d’éthique Paris Nord N°121” and the “Ministère de l’enseignement supérieur, de la recherche et de l’innovation” under the authorization number APAFIS#25642. Basoph8x4C13R(Pellefigues *et al*., 2019) mice were bred on a pure C57BL/6J background in specific pathogen-free conditions at the Malaghan Institute of Medical Research Biomedical Research Unit. All experimental protocols on Basoph8x4C13R(Pellefigues *et al*., 2019) mice were approved by the Victoria University of Wellington Animal Ethic Committee (Permit 24432). Each experiment was performed according to institutional guidelines. MCPT8^DTR^ basophil depletion was induced by intraperitoneal injection of 1µg of diphteria toxin (“DT”, unnicked from Sigma-Aldricht) at the specified times. Blood was harvested by retroorbital puncture on live animals or intracardiac puncture after euthanasia. “Aged” mice refer to animals between 75 weeks and 90 weeks old while “young” animals refer to 8 to 14 weeks old animals, unless specified otherwise (Figure 8). Animals were anesthetized using 3% Isoflurane in an anesthesia system (MiniHub, TemSega) for both wounding and blood harvesting on live animals. Euthanasia was always performed in a controlled CO2 chamber (TemSega). In the “Ear punch” model, a sterile punch biopsy of 2mm was used at the center of the ear to create a circular wound (Kai medical). In the “Ear Tag” model, we used a 2mm plier-style ear punch (Fisherbrand, Thermo Fischer) to generate a circular wound at the center of the ear.

### Tissue pathology and microscopy

Skin Formalin-fixed paraffined embedded (FFPE) 4µm sections of the wounds were stained with Masson’s trichrome to quantify epidermal thickness and collagen deposition using ImageJ v1.54h (Fiji, NIH). For whole-mount immunofluorescence, wholemount ear leaflets were stained with SYTO9 (1/50000) for nuclear counterstaining in Superblock (both from Thermo Fischer) for tdTomato basophils quantification using the CT-M8 mice. Alternatively, ear leaflets of C57BL/6J were permeabilized 10’ in 0.3% Triton X100 (Sigma) in PBS, rinsed in PBS, and then stained 1h in PBS 2% BSA + 10µg/mL 2.4G2 with anti-vimentin AF647 (Biolegend) and counterstained using 500ng/mL DAPI dilactate (Invitrogen). For neutrophils and monocytes-macrophages quantification, ear leaflets of C57BL/6J were permeabilized 10’ in 0.3% Triton X100 (Sigma) in PBS, rinsed in PBS, and then stained for 1h in PBS 2% BSA + 10µg/mL 2.4G2, 100µg/mL mouse IgG and 100µg/mL rat IgG (Jackson Immunoresearch) in the presence of anti-Ly6G AF488 (1A8) and anti-CD68 AF647 (FA-11) or their isotypes (1/100 each, Biolegend), and counterstained using 500ng/mL DAPI dilactate (Invitrogen), all at room temperature. A maximum projection of 4 to 5 overlapping z-stacks and stitching of 9 fields of view centered around the wound are represented. Immunofluorescence of epidermal keratins was done on FFPE sections of MCPT8^DTR^ mice after Sodium Citrate buffer (10mM Sodium Citrate, 0.05% Tween 20, pH 6.0) antigen retrieval using a steamer (30’), 2 washes in PBS 2’, 1 wash in PBS 0.3% TritonX100 10’, 2 washes in PBS 2’, and then stained overnight at 4C in Superblock (Thermo Fischer) +5% rat serum and the following primary antibodies: rabbit IgG anti-filaggrin (Poly19058) and chicken IgY anti-keratin 14 (Poly9060) from Biolegend and guinea pig IgG anti-keratin 10 from Progen (GP-K10), each at 1/400. For secondary staining, samples are rinsed in PBS 0.05% Tween 20 twice 5’, then stained in Superblock + 5% goat serum (Jackson Immunoresearch) 2h at room temperature using “AffiniPure” donkey F(ab’)2 anti-rabbit IgG AF488, anti-chicken IgY AF594 and F(ab’)2 anti-guinea pig IgG AF647, all at 1/500 from Jackson Imunoresearch. Samples were rinsed twice in PBS 0.05% Tween20 5’, and once in 10mM CuSO4/50mM NH4Cl 10’, and in deionized water before mounting in Shandon Immuno-Mount (Thermo Fischer). Fluorescence was then analyzed on a LSM 780 Airyscan confocal microscope (Zeiss), always at 20x in water immersion. Image analysis was done on ImageJ v1.54h (Fiji, NIH).

### Mouse sample handling and flow cytometry

Immediately after death, cardiac puncture was done using a 25G needle, and a minimum of 700 μL of blood was withdrawn in a heparinized tube. The harvested blood cells were resuspended in 5 mL of ACK lysing buffer (150 mM NH4Cl, 12 mM NaHCO3, 1 mM EDTA, pH 7.4) at room temperature for 3 min, then further incubated for 5 min at 4 °C. Subsequently, 10 mL of PBS were added and the sample was centrifuged at 500G for 5 min. When red blood cells were still present, cells were further incubated in ACK lysing buffer for 5 min at 4 °C and the steps outlined above were repeated until red blood cells were lysed. The remaining white blood cells were resuspended in FACS buffer (“FB”: PBS 1% BSA, 0.05% NaN3, 1 mM EDTA). Ear skin was harvested immediately after sacrifice, split into dorsal and dorsal parts using tweezers, and chopped with scissors in 1mL IMDM (Gibco). Ear skin was then digested for 30’ 37C in a shaking incubator at 150 rpm in 2mL of IMDM containing 1mg/mL Collagenase IV and 200µg/mL DNAse I (Sigma). Digestion was stopped by adding 1mM final EDTA (Gibco) and homogenizing samples at 4C. Samples were then filtered through a 70µm nylon mesh (Falcon).

Cells were then washed in PBS and resuspended in 96 well U-bottomed plates (Costar) before being stained for viability using 1/200 Ghost 510 (Tonbo) 15’ 4 C. Then cells were washed in FB, and stained extracellularly in a blocking solution of FB containing 10µg/mL 2.4G2 (BioXcell) and 100µL/mL of mouse IgG, 100µg/mL rat IgG and 100µg/mL of Armenian Hamster IgG (all from Jackson Immunoresearch) with necessary fluorophore-labeled antibodies for 20’ 4C in the dark (Supplementary Table 1). Then cells were washed twice in FB and fixed in PBS 1% PFA (Sigma-Aldricht) 10’ 4C before being washed in FB, and stored in FB 4C overnight before analysis on a 3 laser Aurora spectral flow cytometer (Cytek, figure 1D, Supplementary Figure 2) or X20 Special Order 5-lasers Fortessa (BD) flow cytometer (other figures). Subsequent analysis was done using FlowJo 10 (BD). Mfi always represents geometrical mean intensity and is used to quantify fluorescence intensity. Alternatively, FSC-A and SSC-A are quantified as median values due to the linear nature of the variable. Results have been normalized as % of change from control to pool data from experiments performed after significant flow cytometer baseline changes.

Skin cytokines or growth factors were quantified using LegendPlex Mouse Cytokine release syndrome and Hematopoietic stem cell kits (Biolegend). Briefly, ear skin was snap-frozen and stored at -80C immediately after euthanasia. Then, tissue was thawed in Cell lysis buffer (Cell Signaling) containing 1/100 Phenylymethanesulfonyl fluoride (Sigma) and minced before being homogenized using 5mm stainless steel beads using a Tissue Lyzer II (Qiagen). Protein content was quantified using a BCA test (Qiagen). Legendplex assay was done as per the manufacturer’s instructions using a X20 Special Order 5-laser Fortessa (BD) flow cytometer.

### Molecular biology

Whole skin gene expression was quantified on ear skin that was snap-frozen immediately after euthanasia and stored at -80C. Then 30 mg of ear skin was used to extract mRNA using the RNeasy fibrous Tissue kit and the Tissue Lizer II, as per the manufacturer’s protocol (Qiagen). cDNA were done using the High-quality RNA to cDNA kit (Thermo Fischer) and stored at -20C until quantitative qPCR was done using the PowerUp SYBR Green Master Mix 2x (Thermo Fisher) using a CFX96 real-time thermocycler (Bio-Rad), as per the manufacturer’s protocol. The primers described in Supplementary Table 2 were used.

### Single-cell RNA-seq data analysis

We reanalyzed our previously published scRNA-seq dataset (Vu et al., 2022) of dorsal wound cells from young (8 weeks old) and aged (88 weeks old) mice at 4 and 7 days post wounding. To identify the basophil cell cluster (MCPT8+ Kit-) and perform differential gene expression analysis of basophil cells between young and aged mice, we performed subclustering analysis of the previously assigned “mast cells” using the Seurat R package (version 4.4.0)(Stuart *et al*., 2019). Alternatively, for visual representation, mast cells and basophils were identified by t-SNE clustering using Cytonaut (https://www.cytonaut-scipio.bio/). To minimize the difference between different versions of Chromium Kits, we only used the cells from v3 runs. The top 10 principal components with a suitable resolution in ‘ FindClusters‘ function were used for identifying the basophil cell cluster. ‘FindAllMarker‘ function with parameters ‘ min.pct ‘ being 0.25 and ‘logfc.threshaged‘ being 0.1 was used to find differentially expressed genes. GO biological process enrichment analysis was performed using the clusterProfiler 4.0 R package(Wu *et al*., 2021). Cell-cell communication analysis was performed as in our previous study using CellChat (version 2.0) (Jin *et al*., 2021; Vu *et al*., 2022; Jin, Plikus and Nie, 2023). Briefly, the communication probability between two interacting cell groups was quantified based on the average expression values of a ligand by one cell group and that of a receptor by another cell group, as well as their cofactors. We computed the average expression of signaling molecules per cell group using 10% truncated mean. The significant interactions were inferred using permutation tests.

### Statistics and data visualization

Statistical analyses were performed and plotted using Prism v9.x or v10.x (GraphPad). Data from experimental replicates were pooled if doing so led to a decreased variance; alternatively, representative results from an individual experiment were represented. Parametric or nonparametric tests were used depending on the normality of the data distribution assessed with a D’Agostino-Pearson K2 omnibus test. Posttest-adjusted p values are always represented when used. A two-tailed p-value less than 0.05 was considered the threshaged for significance. Exact tests and post-tests are always described in the figure’s legend.

## Data availability

Data are available in the article itself and its supplementary materials. The scRNA-seq data was previously reported(Vu *et al*., 2022), and is accessible in the GEO database under accession #GSE188432.

## Authors’ contributions

C.P. designed experiments, conducted experiments, acquired and analyzed data, wrote the manuscript, conceived and directed the project. N.C. and G.L.G. provided funding and edited the manuscript. J.B., V.P., C.S., L.C., O.T., M.K.T., J.T., Q.S., K.N., and G.G. conducted experiments, acquired data, managed animals, and/or edited the manuscript. S.J. analyzed bioinformatic data and/or edited the manuscript. X.D., K.M., H.K., J.X.J., M.B., R. M., and U.B. provided resources and edited the manuscript. C.P. has full access to all of the data in the study and takes responsibility for the integrity of the data and the accuracy of the data analysis. All authors approved the final version of the article.

## Funding

This work was supported by the “Fondation pour la Recherche Médicale” (FRM) (grant # EQU201903007794) and the “Agence Nationale de la Recherche” (ANR) (grants # ANR-19-CE17-0029 BALUMET) to NC and by an independent research organization grant from the Health Research Council of New Zealand and by the Marjorie Barclay Trust to G.L.G. and by the NIH (grant R01 AG045040) to J.X.J.. This research was also funded by the ANR grant ANRPIA-10-LABX-0017 INFLAMEX to the Centre de Recherche sur l’Inflammation, by the “Centre National de la Recherche Scientifique” (CNRS), by “Université de Paris”, and by the “Institut National de la Santé et de la Recherche Médicale” (INSERM).

## Conflicts of interest

The authors declare no conflict of interest with the results of the current study.

## Acknowledgments

We acknowledge the expert work from the members of the UMR1149 “Centre de Recherche sur inflammation” animal (I. Renault and S. Olivré), flow cytometry (J. Da Silva, and V. Gratio), and imaging core facilities (S. Benadda); and the help from L. Wingertsmann from the morphology core facility (INSERM UMR1152). We also wish to thank the expert support of the Malaghan Institute of Medical Research Hugh Green Cytometry Core, Research Information Technologies, and Biomedical Research Unit staff.

## Notes

### Competing Interest Statement

The authors have declared no competing interest.

