## Supplementary Figures for "Basophils drive the resolution and promote wound healing in adult and aged mice"

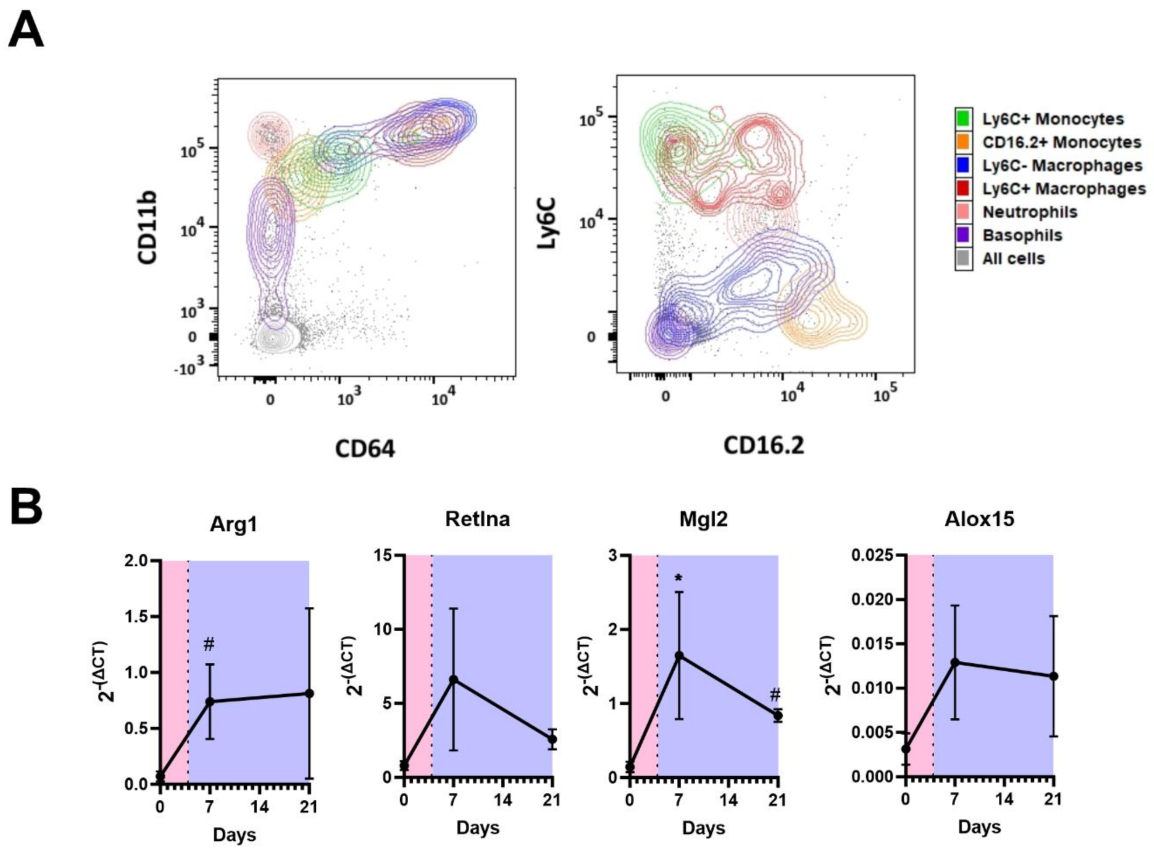


**<u>Supplementary Figure 1: Phenotype of skin leukocytes and expression of M2 macrophage-associated genes during the resolution of skin wound healing</u>**. Wild-type mice were wounded by a sterile 2mm punch biopsy of the ear at day 0. A) Representative contour plots showing the expression of CD11b, CD64, Ly6C, and CD16.2 by various skin leukocytes, as observed by flow cytometry at D7 post-wound. B) Gene expression of selected genes was quantified over time after normalization on GAPDH expression (n=4-5). Results are pooled from 3 independent experiments. The inflammation and resolution phases are represented in red or blue with a D7 cutoff. Statistics are a Kruskall-Wallis test corrected by a Dunn's multiple comparison test against day 0. $p<0.01$: #; $p<0.05$: *

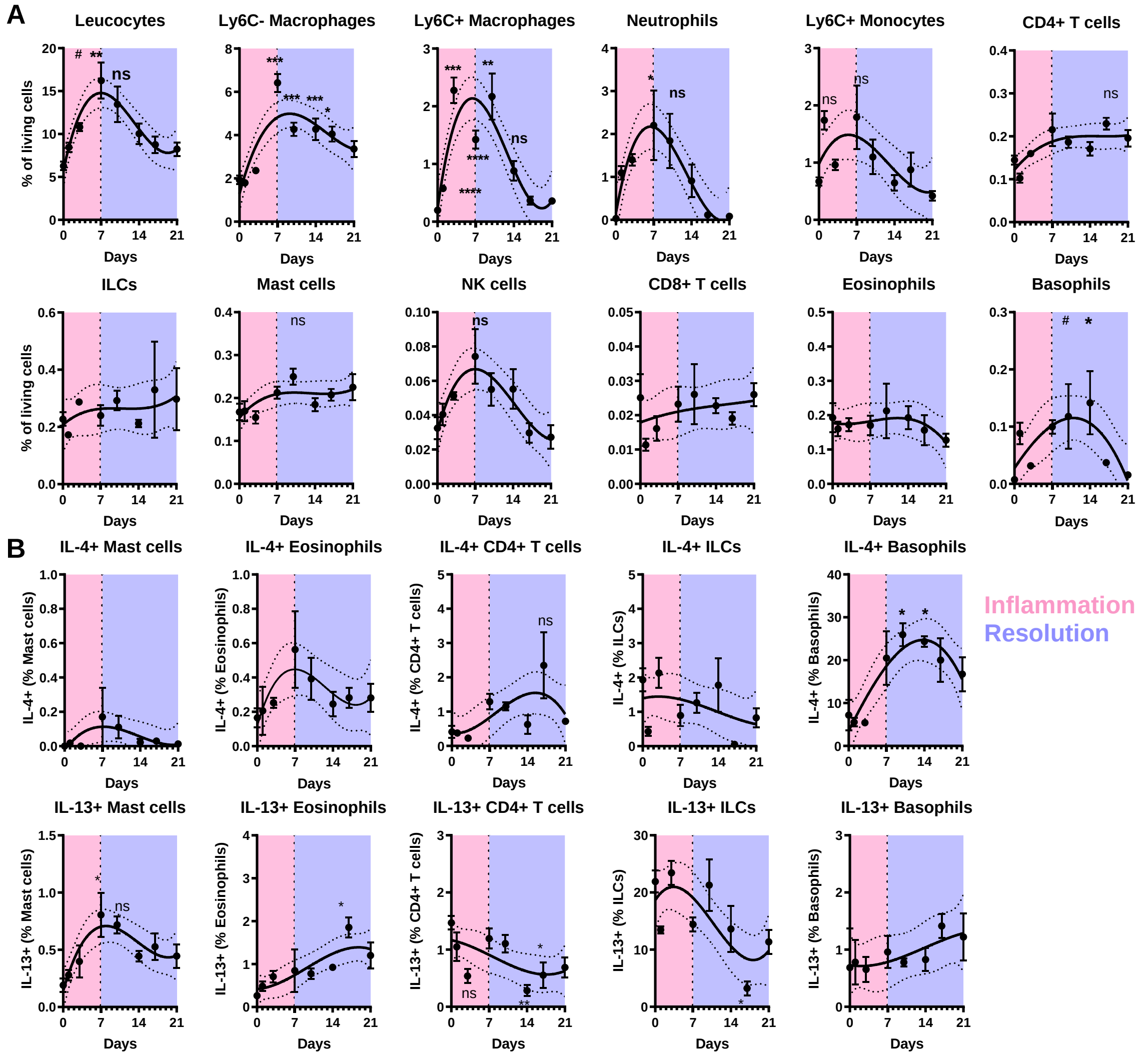


**<u>Supplementary Figure 2</u> Kinetics of leukocyte infiltration and Type 2 cytokines single cell expression in ear skin wounds**. A) Leukocyte infiltration was analyzed by spectral flow cytometry over time after a 2mm sterile punch, scissor-type in Basoph8x4C13R mice, and B) their expression of IL-4 and IL-13 was quantified (n=4). Results are from a single experiment. The inflammation and resolution phases are represented in red or blue with a day 7 cutoff, as from the peak of leukocytes and neutrophils infiltration. A 3rd order polynomial interpolation with a 95% confidence interval is represented. Statistics are ordinary one-way ANOVA corrected by a Holm-Sidak's multiple comparison test against day 0. p<0.01: #; p<0.05: *; p<0.01: **; p<0.001: ***; p<0.0001: ****

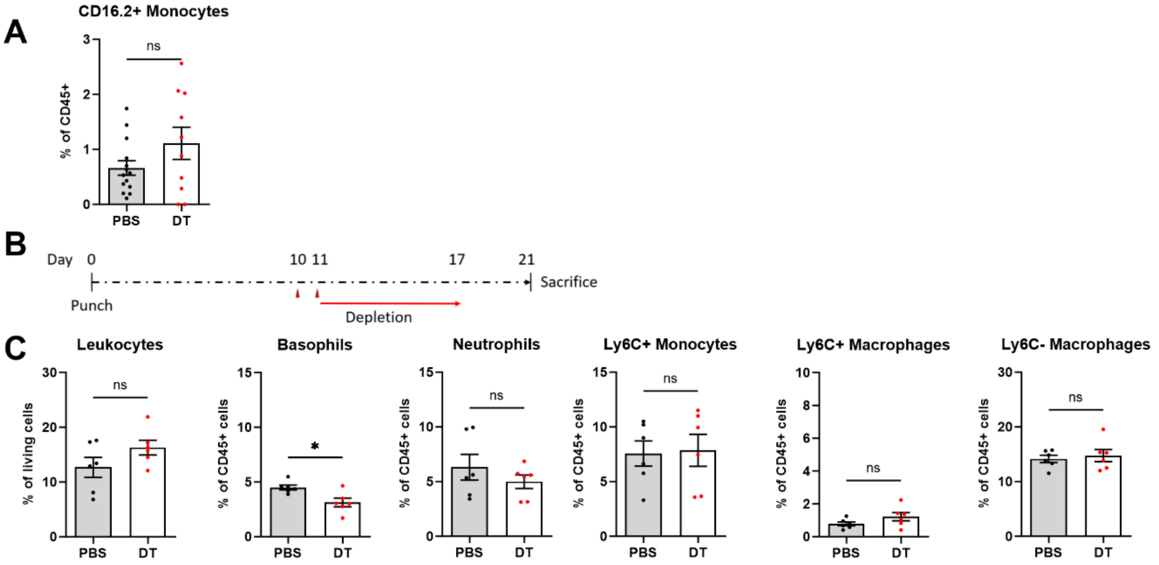


**<u>Supplementary Figure 3: Effects of basophils transient depletion on the skin wound environment.</u>** A) MCPT8$^{DTR}$ mice were ear punched and injected with diphteria toxin (DT) or PBS at Days 13, 14, and 19 before being sacrificed on Day 21. The proportion of their CD16.2+ monocytes was quantified by flow cytometry (n=14, 10). B) Alternatively, MCPT8$^{DTR}$ mice were ear punched and injected with diphteria toxin (DT) or PBS on Days 10 and 11 before being sacrificed on Day 21. C) Various leukocyte subsets were then quantified in the skin (n=6). Results are A) pooled from two to three independent experiments, or C) come from a single experiment. Statistics are two-tailed Mann-Whitney tests. ns: non significant; $p<0.05$: *

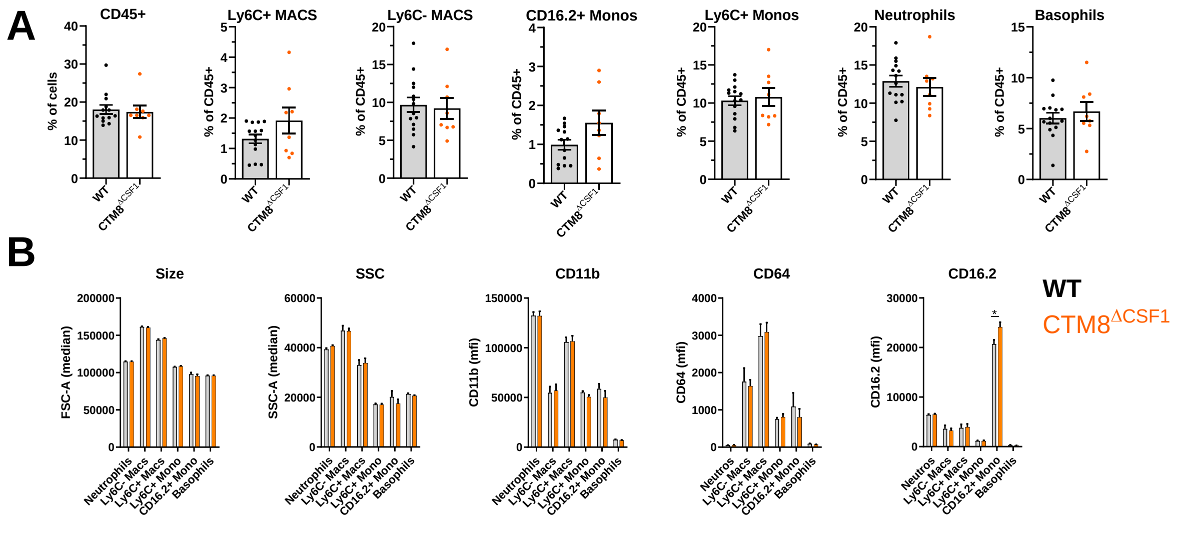


**Supplementary Figure 4: Basophil-derived CSF1 shows minimal immunoregulatory properties at day 7 post-wound.** Wild-type (WT) and CTM8$^{\Delta CSF1}$ littermates were ear-punched and sacrificed 7 days later. A) the proportion of skin wound leukocytes and B) their phenotype was analyzed by flow cytometry (n=14, 8). Results come from a single experiment. Statistics are two-tailed Mann-Whitney tests. $p<0.05$: *

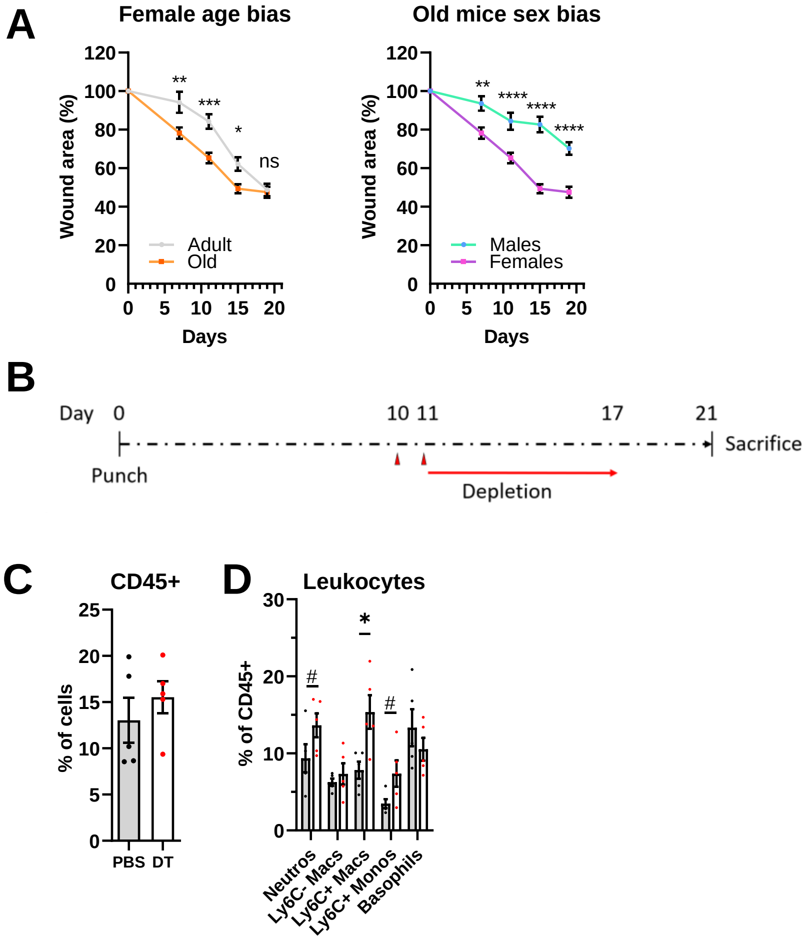


**Supplementary Figure 5: Age and sex related biases in wound closure rate and transient depletion of basophils during the resolution phase of old mice wound healing.** A) Adult and old female mice (n=14), and old males and females (n=14), bred in specific pathogen free conditions, were ear punched and the rate or their wound closure was monitored overtime. B) Old MCPT8-DTR mice were ear punched at D0 and depleted of basophils by injections of diphteria toxin at D10 and D11, before being sacrificed at D21. Their ear skin was then analyzed for C) total leukocyte content and D) specific immune cells proportions by flow cytometry (n=5). Statistics are A) two-way ANOVAs with a Holm-Sidak's post test comparing both groups at each time point, or C, D) two-tailed Mann Whitney's tests. p<0.1: #; p<0.05: *; p<0.01: **; p<0.001: ***; p<0.0001: ****
