## Supplementary Table 1 for "Basophils drive the resolution and promote wound healing in adult and aged mice"

| Clone | Target | Fluorochrome | Source |
|---|---|---|---|
| 2B8 | CD117 | APC/Fire 750 | Biolegend |
| 2B8 | CD117 | BB700 | BD |
| M1/70 | CD11b | BV605 | Biolegend |
| M1/70 | CD11b | BUV737 | BD |
| M1/70 | CD11b | PE-Cy7 | Biolegend |
| 5B11 | CD123 | PE | Biolegend |
| 9E9 | CD16.2 | BV785 | Biolegend |
| 9E9 | CD16.2 | AF647 | Biolegend |
| OX110 | CD200R | PerCPeF710 | ThermoFischer |
| Ba13 | CD200R3 | PE | Biolegend |
| Ba13 | CD200R3 | APC | Biolegend |
| C068C2 | CD206 | AF647 | Biolegend |
| C068C2 | CD206 | PE-Cy7 | Biolegend |
| PC61 | CD25 | BB515 | BD |
| TY25 | CD273 | BV421 | Biolegend |
| TY25 | CD273 | APC | Biolegend |
| 145-2C11 | CD3 | PeCy5 | Thermofischer |
| 145-2C11 | CD3 | BV605 | Biolegend |
| URA-1 | CD301b | PerCP/Cyanine5.5 | Biolegend |
| RM4-5 | CD4 | AF647 | Biolegend |
| RM4-5 | CD4 | BV605 | BD |
| 30-F11 | CD45 | BUV395 | BD |
| 30-F11 | CD45 | APC-Cy7 | Biolegend |
| 30-F11 | CD45 | APC-Cy5,5 | Antibodies-online.com |
| DX5 | CD49b | FITC | Biolegend |
| X54-5/7.1 | CD64 | Pe-Dazzle 594 | Biolegend |
| X54-5/7.1 | CD64 | BV421 | Biolegend |
| FA-11 | CD68 | AF647 | Biolegend |
| 53-6.7 | CD8a | PerCPCy55 | BD |
| 53-6.7 | CD8a | BV785 | BD |
| MAR-1 | FcεRIα | AF647 | Biolegend |
| MAR-1 | FcεRIα | FITC | Biolegend |
| M5/114.15.2 | I-A/I-E | Pacific Blue | Biolegend |
| RME-1 | IgE | FITC | Biolegend |
| R6-60.2 | IgM | PE-Cy7 | Biolegend |
| HK1.4 | Ly-6C | AF700 | Biolegend |
| HK1.4 | Ly-6C | BV785 | Biolegend |
| 1A8 | Ly-6G | FITC | Biolegend |
| 1A8 | Ly-6G | BV650 | Biolegend |
| PK136 | NK1.1 | BV650 | BD |
| E50-2440 | SiglecF | Pe | Biolegend |
| 53-2.1 | CD90.2 | PeCy7 | BD |
