## Supplementary Table 2 for "Basophils drive the resolution and promote wound healing in adult and aged mice"

| oligo name | sequence 5' | Company |
|---|---|---|
| alox15 FOR | CGGCGACCAGTATCTCTGAC | Life |
| alox15 REV | TTCCAGGAGTTTCGAACCCG | Life |
| Arg1 FOR | GTGAAGAACCCACGGTCTGT | Life |
| Arg1 REV | AGAAAGGACACAGGTTGCCC | Life |
| mgl2 FOR | GGAGTCTCCAAAGTTTGCTCT | Life |
| mgl2 REV | GGTGCCTAGGTCCCTCCTTA | Life |
| retnla FOR | ACTATGAACAGATGGGCCTCC | Life |
| retnla REV | TCCACTCTGGATCTCCCAAGA | Life |
| il4 FOR | CTGGATTCATCGATAAGCTG | Sigma |
| il4 REV | TTTGCATGATGCTCTTTAGG | Sigma |
| csf1 FOR | TAGAAAGGATTCTATGCTG | Sigma |
| csf1 REV | CTCTTTGGTTGAGAGTCTA | Sigma |
| Gapdh FOR | AATGGTGAAGGTCGGTGTGA | Sigma |
| Gapdh REV | GCAACAATCTCCACTTTGCCA | Sigma |
